# The *Yersinia pestis* effector YopM binds the human pyrin death domain to inhibit inflammasome activation and effector-triggered immunity

**DOI:** 10.64898/2026.02.25.707193

**Authors:** Bethany W. Mwaura, Adam R. Simard, Kate M. Brown, Morgan L. Pettis, Ao Liu, Dean R. Madden, James B. Bliska

## Abstract

Bacteria in the genus *Yersinia* use a type III secretion system to inject several effectors into host cells to disrupt multiple signaling processes. Two effectors disrupt host signaling by inactivating RhoA GTPases. Inactivation of RhoA triggers pyrin inflammasome assembly in infected phagocytes leading to a protective immune response. A third protein, YopM, is an essential virulence factor that counteracts effector-triggered immunity by inactivating pyrin. YopM has a leucine-rich repeat (LRR) domain and hijacks host kinases to inactivate pyrin by phosphorylation. Previously, the LRR domain of *Yersinia pestis* YopM was implicated in binding to an N-terminal region of human pyrin, which includes the eponymous pyrin domain (PYD), a member of the death domain family. However, the interaction mechanism was undefined. Using a bacterial two-hybrid assay, protein biochemistry and X-ray crystallography, we determined that the concave surface of the YopM LRR domain binds to the PYD. Identification of critical residues in the interaction revealed that YopM binds PYD R42, an amino acid important for positive and negative regulation of pyrin. YopM binding to the PYD was reduced by R42W, a codon change associated with the autoinflammatory disease Familial Mediterranean Fever (FMF). Furthermore, we show that YopM codon substitution variants defective for PYD binding fail to inhibit the pyrin inflammasome in human monocytes infected with *Yersinia*. In addition, we found that PYD binding is dispensable for YopM to inhibit the inflammasome in murine macrophages, suggesting this effector uses a distinct mechanism to target mouse pyrin. These results define how *Y. pestis* YopM binds the human PYD and provide insights into how this interaction likely selected for pyrin gain of function variants resulting in FMF.

**SIGNIFICANCE:** Pyrin is an inflammasome sensor encoded by the *MEFV* gene that is notable for its role in the autoinflammatory disease Familial Mediterranean Fever (FMF) and immunity to the plague agent *Yersinia pestis*. *Y. pestis* normally prevents inflammasome activation using the virulence factor YopM which binds pyrin and inhibits effector-triggered immunity. Gain of function mutations in *MEFV* that cause FMF were likely selected during historic plague pandemics to counteract YopM. Here, we defined the molecular mechanism of the YopM-pyrin interaction, which revealed that the effector binds to a key site of positive and negative regulation on the N-terminal death domain. These findings have important implications for understanding how YopM promotes pathogenesis and likely selected for gain of function mutations in *MEFV*.

## INTRODUCTION

Pathogens use a variety of strategies to evade detection and elimination by host immune responses. One method employed by numerous pathogenic bacteria involves injecting effector proteins directly into eukaryotic cells. These effectors can be injected via the type III secretion system (T3SS) which is employed by Gram-negative bacteria that infect both animals and plants (1). The T3SS is a sophisticated protein secretion machinery which spans from the bacterial membrane through the host plasma membrane (2). Human pathogenic species of *Yersinia*, *Shigella* and *Salmonella* among others use a contact-dependent T3SS to secrete effectors, regulate host cell responses and promote pathogenesis (2). Host cells can sense assault by the T3SS and or effectors and mount immune responses (3). In macrophages infected with bacterial pathogens, perturbations induced by T3SSs and/or effectors can result in the assembly of inflammasomes (4). Canonical inflammasomes are multiprotein assemblies that promote the recruitment and maturation of caspase-1 into its active form (4). Active caspase-1 cleaves gasdermin-D (GSDMD), pro-IL-18 and pro-IL-1β (5, 6). The N-terminal domain of GSDMD forms pores in the plasma membrane, promoting release of mature IL-18 and IL-1β (5, 6). GSDMD pores can also result in the release of additional pro-inflammatory molecules and intracellular bacteria as a result of cell death, termed pyroptosis (6, 7). Pyroptosis downstream of GSDMD pore formation requires plasma membrane rupture mediated by the ninjurin 1 protein (8). To counteract protective immune responses mediated by IL-1β, IL-18 and pyroptosis, virulent bacterial pathogens have evolved the ability to inhibit caspase-1 inflammasomes in host cells (4, 9).

The genus *Yersinia* contains several Gram-negative bacterial pathogens that cause invasive human infections ranging in severity from self-limiting mesenteric lymphadenitis (*Y. pseudotuberculosis*) to lethal plague (*Y. pestis*). *Yersinia* colonize lymphoid tissues and replicate as extracellular microcolonies in phagocyte-rich lesions known as pyogranulomas and a plasmid-encoded T3SS is required for this infection process (10, 11). The T3SS delivers seven effectors named *Yersinia* outer proteins (Yops) into the cytosol of host cells in contact with the bacteria (12). The effectors inhibit key host cell processes, such as proinflammatory cytokine production and phagocytosis (12). YopE and YopT are two *Yersinia* effectors that contribute to the anti-signaling functions of the T3SS by targeting Rho GTPases (13, 14). YopE has a GAP domain that accelerates conversion of GTP to GDP, locking the GTPase in an inactive form (13). YopT is a cysteine protease which cleaves off a C-terminal amino acid and lipid anchor from the GTPase, disrupting its association on host membranes and preventing interaction with downstream targets (14). YopM is a leucine rich repeat (LRR) effector that inhibits the pyrin inflammasome in activated macrophages (15, 16). YopJ is a member of the acetyltransferase effector family (17) that inhibits MAPK and NF-kB signaling pathways to inhibit proinflammatory cytokine production (18, 19). YopJ also cooperates by an unknown mechanism with YopM to inhibit the pyrin inflammasome in activated macrophages (20).

Inactivation of Rho GTPases (RhoA, –B or –C, hereafter referred to as RhoA) by bacterial toxins or effectors triggers assembly of the pyrin inflammasome by an indirect guard mechanism in phagocytes (21, 22). YopE or YopT triggers assembly of the pyrin inflammasome in activated macrophages infected by *Yersinia* (15, 16, 23, 24). Pyrin, encoded by the Mediterranean fever (*MEFV*) gene, is a cytosolic pattern recognition receptor (PRR) which is preferentially expressed in phagocytes such as macrophages and neutrophils (25, 26). Human pyrin is comprised of the following domains: An N-terminal PYD followed by a linker region, B-box domain, coiled coil domain (CC) and a C-terminal B30.2 domain (Fig. S1A). Murine pyrin is structurally like human pyrin except that it lacks the C-terminal B30.2 domain, which is replaced by a short unstructured tail (Fig. S1A) (23, 25, 26). Pyrin is also known as TRIM20 as it belongs to the TRIM family of proteins (27) and forms dimers via the CC domain (Fig. S1B) (28). By analogy to TRIM5 (29), the pyrin B-box potentially mediates the formation of higher order oligomers such as dimers of dimers (30). Pyrin is negatively and positively regulated by multiple mechanisms (21, 23, 30–35). Pyrin is negatively regulated by phosphorylation via the protein kinase C-related kinase (PRK, also known as PKN, protein kinase N) which in turn is activated by RhoA. Human pyrin is phosphorylated on serine residues 208 and 242 (205 and 241 on mouse pyrin) on the linker region creating a binding site for 14-3-3 protein dimers (32, 33). Pyrin is also negatively regulated by binding of the PYD to the B-box (31). A model posits that phosphorylated pyrin is rendered inactive as the 14-3-3-bound linker is bent into a hairpin (36) and the PYD is sequestered by the B-box (Fig. S1B), preventing interaction with apoptosis-associated speck-like protein containing a CARD (ASC) (23). In *Yersinia-*infected cells pyrin should be activated when YopE or YopT inactivate RhoA, and a positive regulation step leads to dephosphorylation by a phosphoprotein phosphatase (23, 24, 30). According to the model, when pyrin is dephosphorylated, the PYD is released from the B-box and exposed, ASC is recruited, caspase-1 is activated, GSDMD pores are formed and active IL-1β and IL-18 are secreted (23, 30). However, pyrin inflammasome activation in phagocytes infected with *Yersinia* is blocked due to the activity of YopM (15, 16). YopM hijacks the host kinases PRK and ribosomal S6 kinase (RSK) (37, 38) to phosphorylate pyrin, locking it in an inactive state (15, 23, 39). YopM is crucial for *Yersinia* virulence in mouse models of infection (16, 40, 41) and yet a *Y. pseudotuberculosis ΔyopM* mutant is fully virulent in *Mefv*^-/-^ mice, demonstrating that pyrin is the major host target of YopM (15).

YopM is a member of the bacterial subfamily of leucine rich repeat (LRR) effector proteins (42). Pathogenic species and strains of *Yersinia* encode for a variety of different YopM proteins, referred to here as isoforms, with varying numbers of LRRs. Despite these differences in the number and composition of LRRs, all YopM isoforms studied so far have been shown to inhibit the pyrin inflammasome (15, 16). The structure of YopM from *Y. pestis* 195/P has been determined and adopts an overall horse-shoe shaped fold comprised of two N-terminal α-helices followed by an extended LRR region with short β-strands (43). The two α-helices provide a nucleation site that facilitates the folding of the LRRs (43). The N– and C-termini of YopM are unstructured and yet highly conserved among YopM isoforms. The N-terminus contains the secretion signal required for YopM translocation into host cells (44), and the C-terminal tail is a binding site for the N-terminal kinase domain of host kinase RSK (Fig. S1A) (40, 41, 45).

During *Yersinia* infection of host phagocytes, YopM acts as a scaffold connecting the host kinases RSK and PRK (37, 38) to allow for pyrin phosphorylation and inactivation (15, 16, 39). Binding to YopM results in autophosphorylation and activation of RSK, which in turn phosphorylates and activates PRK (Fig. S1C) (37). Inflammasome activation was increased when murine macrophages infected with *Y. pseudotuberculosis* were treated with a PRK inhibitor or with siRNAs that reduced expression of *Prk*, suggesting that PRK activity is important for YopM to inhibit pyrin in mouse phagocytes (15). Similarly, inflammasome activation was increased when human monocytes infected with *Y. pestis* were treated with RSK inhibitors or with siRNAs that reduced expression of *Rsk*, suggesting that RSK activity is important for YopM to inhibit pyrin in human phagocytes (39). Park *et al*. used *in vitro* kinase assays with purified proteins to show that *Y. pestis* YopM could hijack either PRK or RSK to phosphorylate the human pyrin linker, but RSK had much higher activity, and there was no additive effect when both kinases were present (39). These data suggest that both kinases are important, but RSK may be critical for *Y. pestis* YopM to inhibit human pyrin.

Like other proteins in this family, the concave surface of the YopM LRR domain contains exposed amino-acid sidechains (43) and is likely important for its scaffolding function in binding host proteins. The kinase domain of PRK binds to YopM (Fig. S1A) (37), and the interaction site appears to be within LRRs 6-15 of the *Y. pestis* isoform (40). Deletion analysis with purified *Y. pestis* YopM variants demonstrated an importance for LRRs 1 and 2 for pyrin binding (39). However, a previous study showed low expression in *Yersinia* of a YopM Λ1LRR2-3 variant (41), suggesting that this deletion disrupted YopM stability. Therefore, it’s possible that the reduced binding of YopM ΔLRR 1-2 to pyrin (39) was a result of structural instability caused by the deletion, and the pyrin binding site on YopM is yet to be established. Additionally, it has been shown that YopM binds to the N-terminus of human pyrin encompassing the PYD and linker (39). Removal of the 90 amino acid PYD prevented phosphorylation of human pyrin by YopM-hijacked RSK in an *in vitro* kinase reaction (39), revealing a key role for this N-terminal domain, but the molecular mechanism underscoring this interaction is not fully understood.

Here, we provide evidence of direct binding between *Y. pestis* YopM isoforms and the human PYD. We determined two co-crystal structures of YopM bound to human PYD and identified key residues on each that are involved in their interaction. We also identified YopM variants with alanine substitutions in LRR 5 that are defective for PYD binding *in vitro* and used human monocyte infection assays to demonstrate that PYD binding is critical for YopM to target and inhibit the pyrin inflammasome *in vivo*. Our data also indicate that *Y. pestis* YopM specifically binds a key regulatory site in the PYD, which has implications for the evolutionary host-pathogen arms race that resulted in FMF (39) and may explain why a distinct mechanism appears to be used to target the mouse pyrin inflammasome.

## RESULTS

### YopM Binds to the PYD of Human Pyrin

We tested for direct interactions between *Y. pestis* YopM isoforms and N-terminal domains of human or murine pyrin. We first used the bacterial adenylate cyclase-based two hybrid assay (BACTH) to test for protein-protein interactions. Two different *Y. pestis* YopM isoforms were studied, the 13-LRR version from Pestoides A (YopM^PestA^) and the 15-LRR version from KIM (YopM^KIM^) (46), which have been shown to inhibit the pyrin inflammasome (15, 16). YopM^PestA^ and YopM^KIM^ were expressed as N-terminal fusions with the T25 domain and the human (h) or murine (m) PYD domains as C-terminal fusions to T18 in *Escherichia coli*. The hPYD and mPYD share 76% identity at the primary sequence level. YopM^PestA^ and YopM^KIM^ showed similar strong binding to hPYD (Fig. 1A, Fig. S2A).

**Fig. 1.**
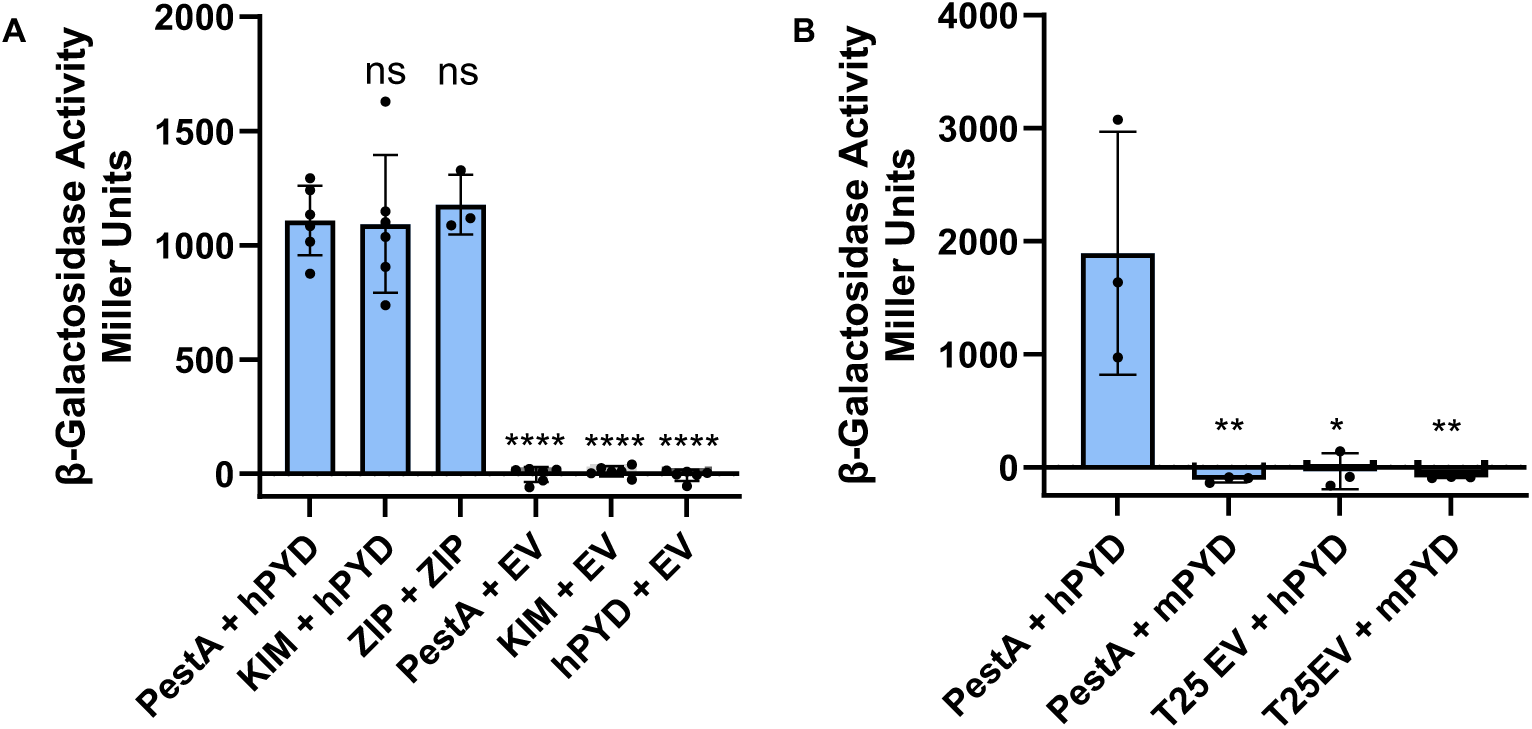
Analysis of YopM-PYD interactions using BACTH. The indicated YopM proteins were N-terminally tagged with T25 and hPYDs (**A, B**) or mPYDs (**B**) were C-terminally tagged with T18. Plasmids encoding the fused partners, ZIP positive controls or empty vector (EV) controls were co-transformed into *E. coli* to test for protein-protein interactions. Measured Miller units of β-galactosidase activity indicating protein interaction in *E. coli* are indicated on the y-axis. Results are averages and standard deviations with data pooled from at least 3 independent experiments. Significance determined by one-way ANOVA and Tukey’s post hoc test in comparison to PestA + hPYD.****p<0.0001; **p=0.01; *p=0.05; ns, not significant.

Surprisingly, YopM^PestA^ showed no detectable binding to mPYD (Fig. 1B). We further investigated these interactions using GST pulldowns, whereby GST-tagged YopM^KIM^ was co-expressed in *E. coli* with His-tagged mPYD or hPYD. Samples of clarified *E. coli* input lysates or pull downs were analyzed by SDS-PAGE and blue staining or anti-His immunoblotting. As compared to the GST-alone control, there was low but detectable pull down of mPYD with GST-YopM^KIM^ (Fig. S3). However, as compared to mPYD, there was increased soluble hPYD in the input lysates, and as a result, more hPYD was pulled down with GST-YopM^KIM^ (Fig. S3). Overall, these results suggest that YopM^PestA^ and YopM^KIM^ strongly bind to hPYD, increasing its solubility, and have a lower affinity for mPYD (Fig. S1A).

Next, we followed up on previous experiments that identified the human pyrin linker as a YopM^KIM^ binding site (39). We tested a fragment of the human pyrin linker (amino acids 186-326) which was shown to bind to YopM^KIM^ in GST pull-downs (39). However, we found that this linker fragment (hPyrin186-326) had non-specific binding to GST (Fig. S4A). In addition, we found that YopM^KIM^ did not bind to hPyrin186-326 in BACTH (Fig. S4B). In summary, these results indicate that the PYD is the primary targeting domain for *Y. pestis* YopM isoforms in the human pyrin N-terminal region (Fig. S1A). Since YopM^KIM^ and YopM^PestA^ inhibit mouse pyrin in macrophages and *in vivo* (15, 39), there may be a different mechanism of inflammasome targeting by this effector in murine macrophages, including possible multidentate interactions with the PYD and the linker or other domains, or the use of hijacked PRK as an adapter.

### In-Solution State of YopM^PestA^

A crystal structure of the 15-LRR YopM protein from *Yersinia pestis* 195/P has been determined (43). In that study, the authors found that YopM^195/P^ formed superhelical tetramers in the crystal lattice. The biological relevance of YopM tetramers is unknown, and it was hypothesized to be a crystallographic artifact given that a tetrameric state was not observed in solution (43). However, other studies have shown that YopM forms higher-order oligomers when analyzed by native PAGE in purified form (dimers and tetramers) (41) or in lysates of *Yersinia*-infected macrophages (30, 41), and in the latter case YopM also appeared to interact with dimers and higher-order multimers of the host target protein pyrin (30). We ultimately aimed to determine the structure of the smaller 13-LRR YopM^PestA^ isoform in complex with the hPYD, but first sought to determine the oligomeric state of YopM^PestA^ in solution. We performed analytical size exclusion chromatography (SEC) of purified full-length YopM^PestA^. We observed that YopM^PestA^ elutes at a faster rate than expected for a monomer on SEC compared to globular standard proteins, implying that YopM^PestA^ could be a dimer in solution (∼90kDa) (Fig. S5A,B), consistent with prior work. Notably, a previous study reported that a 20-LRR YopM from *Y. enterocolitica* was a dimer in solution, based on its elution from SEC (47). However, when calibrated against globular standards, SEC retention volume does not accurately predict the molecular mass of non-globular proteins or of proteins which interact with the column matrix (48). Given that YopM is an elongated protein, its migration during SEC may well be unusual compared to the globular protein standards. Therefore, we performed further analysis using SEC coupled with multi-angle light scattering (SEC-MALS) which permits a shape-independent analysis of protein size (48, 49). Data from SEC-MALS indicated that YopM^PestA^ is a monomer (∼42kDa) at neutral pH (7.4) in solution (Fig. 2A). We further utilized velocity analytical ultracentrifugation (vAUC), another shape-independent method of calculating protein size in solution, to analyze purified YopM^PestA^. vAUC data also suggested that YopM^PestA^ is a monomer in neutral pH (Fig. 2B). Interestingly, the sedimentation coefficient of YopM^PestA^ shifted in low pH (5.0) buffer conditions, consistent with a higher molecular mass (∼148 kDa, Fig. 2B). Given that the calculated molecular mass of the trimeric and tetrameric states are ∼126 kDa and ∼168 kDa, respectively, we cannot unambiguously assign an exact stoichiometry for assembly. However, a tetramer is consistent with previously reported crystal structures (43). Taken together with previous observations, our data suggest that YopM^PestA^ is monomeric at neutral pH, and transitions to an oligomeric form under mildly acidic conditions in solution.

**Fig. 2.**
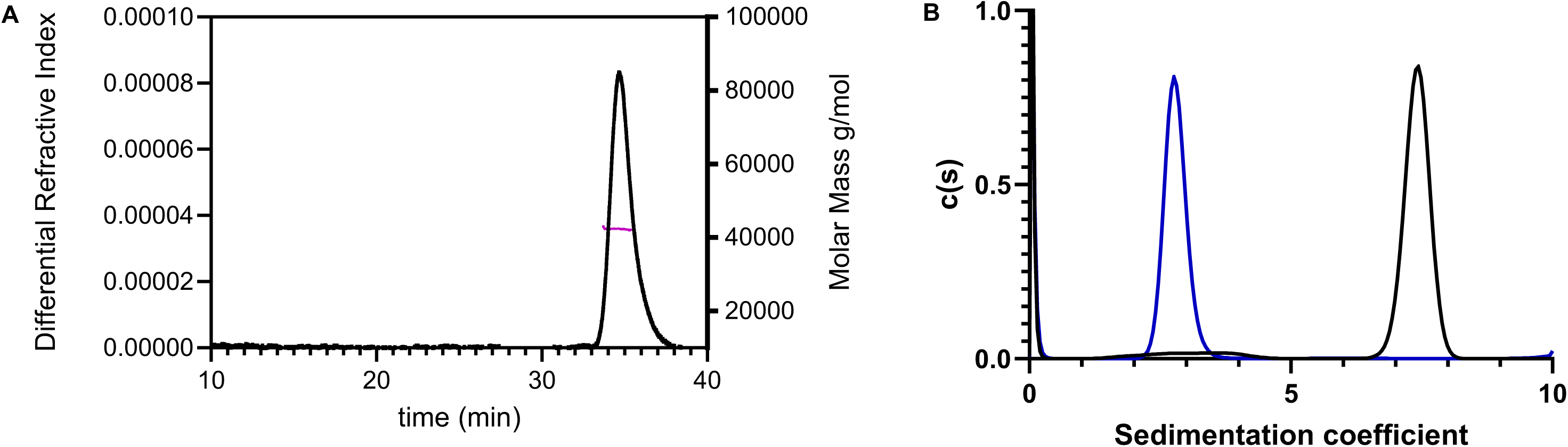
Characterization of YopM^PestA^ in solution by SEC-MALs or vAUC. **A**) SEC-MALS analysis of YopM^PestA^ at pH 7.4. The experimental relative molar mass across the peak width is plotted in magenta (right-hand axis), overlaid with the differential refractive index (left-hand axis) of the chromatogram. The estimated relative molar mass is 42.9 kDa. **B**) Sedimentation coefficient distribution of YopM^PestA^ by vAUC at pH 7.4 (Blue; main peak, corresponding to an estimated M_r_ of 43.5 kDa) and pH 5.0 (Black; main peak, corresponding to an estimated M_r_ of 148.2kDa).

### Structures of YopM^PestA^-hPYD Complexes

Although structural studies of hPYD and closely related YopM isoforms have been conducted in isolation, information pertaining to their complex formation is lacking. Therefore, we sought to determine the structure of a YopM^PestA^-hPYD complex to establish the stereochemical basis for the scaffolding function of YopM. Crystallization trials using the full-length YopM^PestA^ co-purified with hPYD yielded crystals that were difficult to reproduce and optimize (Fig. S6A). Concurrently, a truncation construct of YopM^PestA^ lacking the unstructured first 33 and last 23 residues (43) (amino acids 34-344), was designed to minimize interference with lattice packing by the disordered termini. The resulting protein, henceforth referred to as YopM^ΔNΔC^, has a predicted monomer molecular mass of 35.0 kDa, was produced at high purity, and exhibited excellent biochemical behavior. Importantly, YopM^ΔNΔC^ retained binding to recombinant hPYD as demonstrated by SEC co-elution (Fig. S7). Interestingly, co-elution with hPYD did not significantly alter the SEC retention volume of the complex compared to that of YopM^ΔNΔC^ on its own. In consideration of the aforementioned structural data, we hypothesized that hPYD could ‘tuck’ into the concave surface of YopM^ΔNΔC^ without substantially increasing the hydrodynamic radius of the nascent complex (Fig. S7A,C). Having verified the integrity of the binding interaction, recombinant hPYD was then mixed with YopM^ΔNΔC^ in a 4:1 molar ratio for use in crystallization trials, which led to a condition that produced suitable crystals (Fig. S6B).

In total, X-ray diffraction data were obtained from crystals produced using both strategies and the resulting complexes were found to crystallize in different space groups: YopM^PestA^ co-purified with hPYD crystallized in space group *C* 2, and the YopM^ΔNΔC^-hPYD complex crystallized in space group *P* 3_2_21 (Table 1). Although the former produced quality diffraction to 2.1 Å, it was apparent from the electron density map that the C-terminal portion of YopM^PestA^ beginning in the middle of LRR 9, as well as the sixth α-helix of hPYD, were missing. Careful inspection of the crystal lattice led us to conclude that these missing regions could not be included without significant steric overlap with symmetry-related molecules. Based on these observations, we hypothesize that the fragments captured in this crystal were the result of an unanticipated limited proteolysis event, which is consistent with the difficulties in reproducing this crystal form. This complex is henceforth referred to as YopM^PestA^*-hPYD*. In light of these observations, the co-crystal structure of YopM^ΔNΔC^-hPYD, which diffracted to 3.7 Å, was also refined and used to authenticate the limited proteolysis structure, which possessed the resolution necessary to confidently model the side-chain interactions at the binding interface.

**Table 1.**
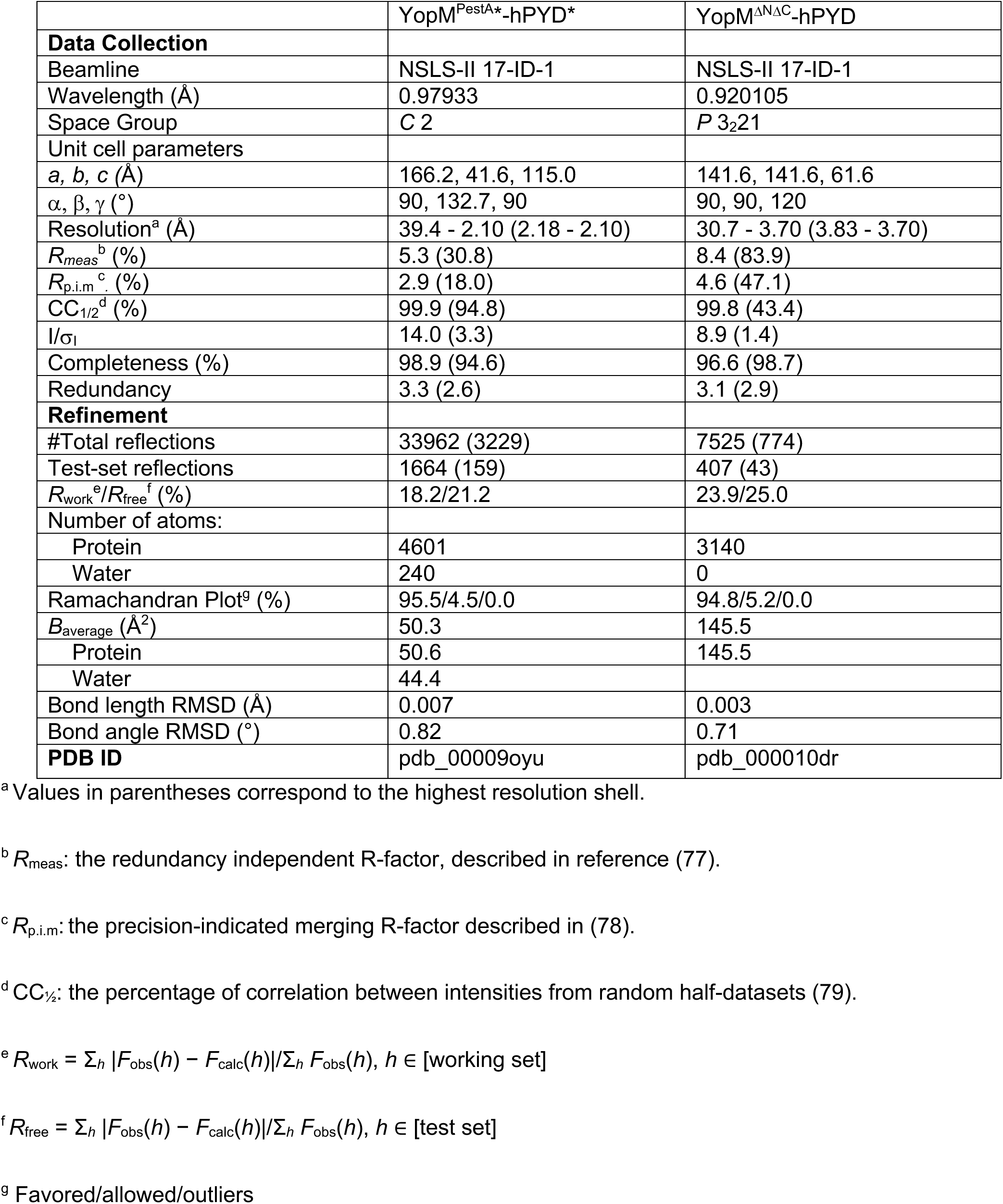
Summary of data collection, reduction, and refinement statistics.

With the exception of the truncations described above, the quaternary structures of both complexes are highly similar. The complex forms in a 1:1 stoichiometry with hPYD* nestled into the concave surface of YopM^PestA^* (Fig. 3A) and is consistent with the SEC co-elution profiles. Overall, the two YopM^PestA^*-hPYD* molecules in the asymmetric unit are similar to each other (C_α_ RMSD = 0.6 Å), but hPYD α2, which lies near the binding interface, was observed in slightly different positions (Fig. S8A). This difference could be attributed to lattice packing, and alignment to the YopM^ΔNΔC^-hPYD complex identified “Molecule 1” as being most similar (compare Fig. S8B, C_α_ RMSD = 0.5 Å to Fig. S8C, C_α_ RMSD = 0.8 Å). Therefore, we conclude that the purported proteolytic truncation bears no influence on complex formation, and the biologically relevant interface is best interpreted from YopM^PestA^*-hPYD* “Molecule 1,” which was used for subsequent analyses as detailed below.

**Fig. 3.**
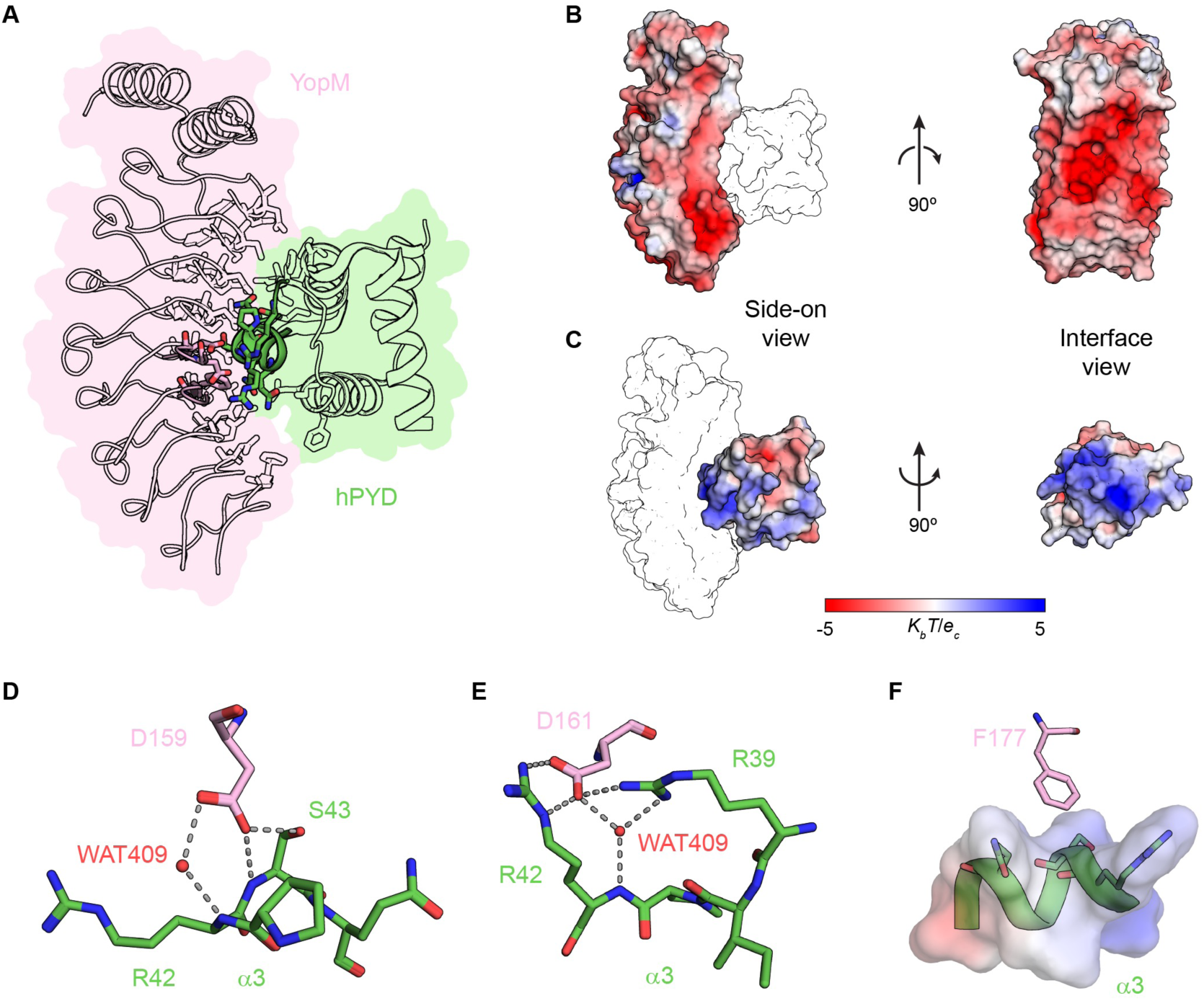
hPYD* binds to the concave surface of YopM^PestA^*. The co-crystal structure of YopM^PestA^*-hPYD* reveals a binding interface composed of the negatively charged concave surface of YopM^PestA^* spanning LRRs 1-9 and the positively charged surface of hPYD* involving α-helices 2-4. **A**) YopM^PestA^* (pink) and hPYD* (green) are depicted as ribbon outlines, with interfacial residues depicted as stick outlines. The YopM^PestA^* residues selected for mutation (YopM Ala2) and cognate hPYD* residues are highlighted as pink and green sticks, respectively. **B and C**) The electrostatic surfaces of YopM^PestA^* (**B**) and hPYD* (**C**) are shown in orthogonal views. As a visual aid, the second protein of the complex is shown as a black-and-white outline in the side-on view (left-hand panel). **D-F**) Focused views of interactions highlighted in (A) between YopM^PestA^* and hPYD* α3 that are disrupted in the YopM Ala2 mutant. YopM^PestA^* D159 (**D**) and D161 (**E**) are the foci of hydrogen-bond networks with hPYD* α3 that involve structural water WAT409 and salt-bridges with hPYD* R39 and R42. YopM^PestA^* F177 (**F**) packs against hPYD* α3, becoming shielded from solvent by apolar and polar atoms of hPYD*. Non-carbon atoms are colored by type: oxygen, red; nitrogen, blue.

The co-crystallized complex reveals an extensive region of van der Waals contact, distributed across YopM^PestA^* LRRs 1-9. The interacting residues of hPYD* are localized to α-helices 2-4, with α3 serving as a focal point (Fig. 3A). Detailed analysis of the contact surfaces suggests that electrostatic attraction is a driving feature of the YopM^PestA^*-hPYD* interaction. The inner, concave surface of YopM^PestA^* carries a uniform, negative charge (Fig. 3B). While the electrostatic surface potential of hPYD* is marked by both positive and negative patches, an equatorial and continuous band of positive charge is found on hPYD* that interfaces with the negatively charged, concave surface of YopM^PestA^* (Fig. 3C). In agreement with a binding interface that favors electrostatic complementarity, several extensive hydrogen-bond networks and salt bridges are observed (Fig. S9).

### Identification of YopM Residues Required for PYD Binding and Inhibition of the Human Pyrin Inflammasome

To identify residues on YopM^PestA^ that are critical for hPYD binding, positions predicted by the complex structure to be important for interaction were subjected to alanine substitution in groups of 3, and the resulting variants were named Ala1-3 (Table 2 and Fig. S10A). The variants were tested for hPYD binding by the BACTH assay, which indicated that Ala2 had significantly diminished binding whereas Ala1 and Ala3 retained binding to PYD at WT levels (Fig. S2B, Fig. S10B, Fig. 4A). Of the positions targeted in Ala2, D159 and D161 are in LRR 5, while F177 is in LRR 6 (Fig. S10A). An analysis of the contact frequency between YopM^PestA^* and hPYD* for each interfacial residue is summarized in Fig. S9. Although each YopM^PestA^ Ala variant contains residues that make substantial contact with hPYD (Fig. S9A,B), YopM^PestA^* D159 and D161 also form a dense electrostatic network with hPYD* α3 that involves multiple hydrogen bonds, two salt bridges, and coordination of a structural water (Fig. 3D,E and Fig. S9B). The interactions made by YopM^PestA^* F177 are entirely different, with the aromatic side chain becoming shielded from bulk solvent by packing against hPYD* α3 during binding (Fig. 3F). We purified the Ala2 variant of YopM^ΔNΔC^ (YopM^ΔNΔC^ Ala2) from *E. coli* at similar yields as YopM^ΔNΔC^. Analysis by SEC showed that YopM^ΔNΔC^ Ala2 eluted similarly to YopM^ΔNΔC^ (Fig. S11A,B vs. Fig. S7A,B). As expected, when incubated at an equimolar ratio with purified hPYD and run on SEC, YopM^ΔNΔC^ Ala2 did not co-elute with hPYD (Fig. S11C,D).

**Fig. 4.**
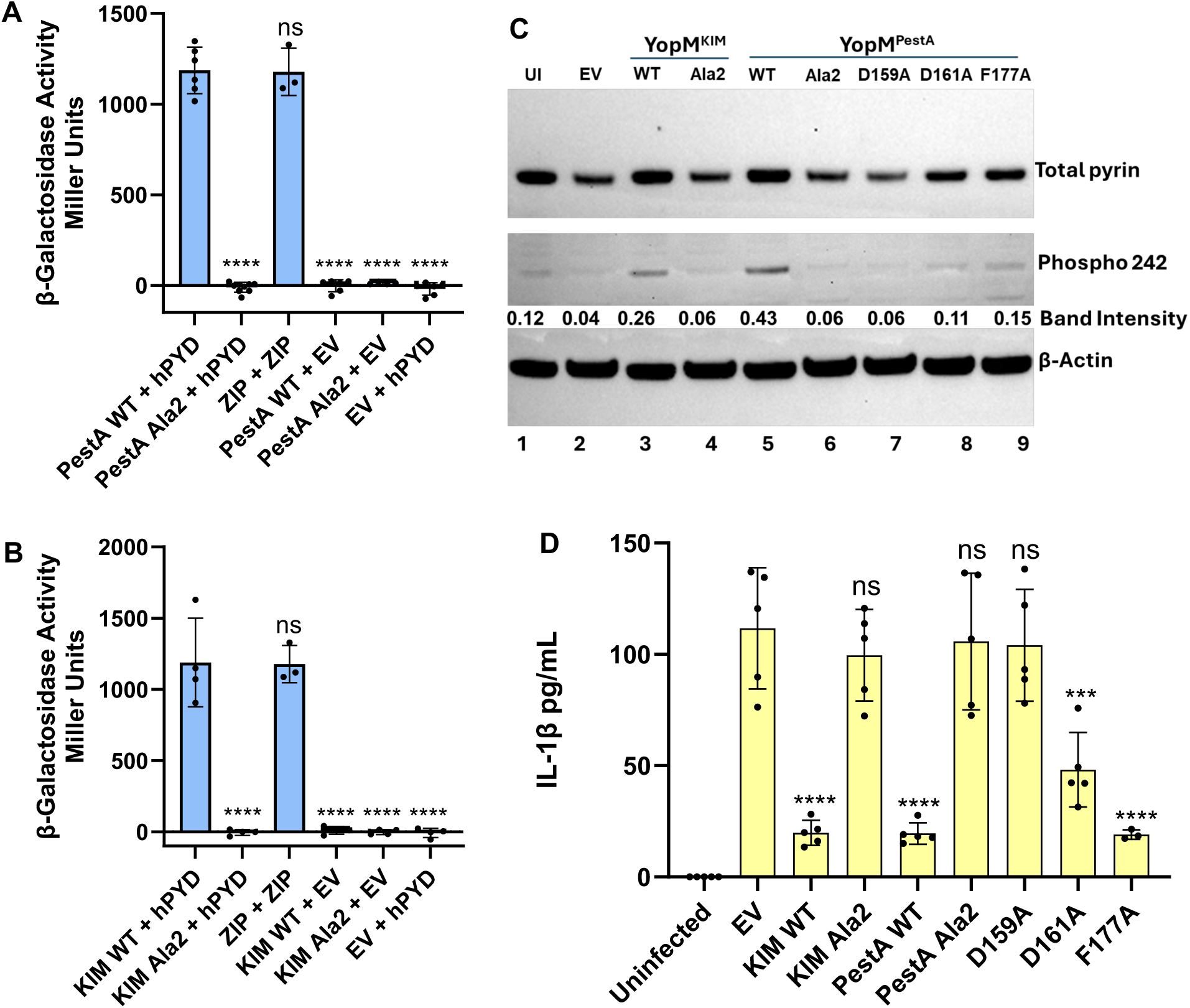
Analysis of YopM variants by BACTH or THP-1 infection assays. **A,B**) The indicated YopM proteins were N-terminally tagged with T25, and hPYDs were C-terminally tagged with T18. Plasmids encoding the fused partners, ZIP positive controls or empty vector (EV) controls were co-transformed into *E. coli* to test for BACTH interactions as in Fig.1. Significance determined by one-way ANOVA and Tukey’s *post hoc* test in comparison to PestA (**A**) or KIM (**B**) + hPYD. **C,D**) THP-1 cells were left uninfected (UI) or infected for 4 h at MOI 30 with *Y. pseudotuberculosis yopJ^C172A^*λ1*yopM* complemented with empty vector (EV) or the indicated YopM isoforms or variants. **C**) Immunoblot analysis of THP-1 lysates with the indicated antibodies. Band intensity signals show the ratio of phospho 242/total pyrin. Results are representative of two independent experiments. **D**) ELISA of IL-1β released from THP-1 cells. Results are averages and standard deviations with data pooled from at least 3 independent experiments. Significance determined by one-way ANOVA and Tukey’s *post hoc* test. Comparisons of each group to EV are shown. ****p<0.0001; ***p<0.001; ns, not significant.

We also tested the impact of the Ala2 D159, D161 and F177 substitutions in the 15-LRR YopM^KIM^ isoform. The sequence alignment of YopM^KIM^ and YopM^PestA^ shown in Fig. S12 indicates that all three positions are conserved. The Ala2 variant of YopM^KIM^ displayed abolished binding to the hPYD in BACTH (Fig. 4B, Fig. S2C).

We next expressed WT or Ala2 variants of YopM^KIM^ and YopM^PestA^ in *Yersinia* for functional studies. A mutant *Y. pseudotuberculosis* strain (*yopJ*^C172A^Δ*yopM*) was complemented with plasmids producing the different YopM proteins under control of native promoters. The resulting strains were tested by Yop secretion assay, which confirmed that the YopM Ala2 variants were released by the T3SS at levels similar to WT, indicating that the amino-acid substitutions did not interfere with protein folding or stability (Fig. S13A lanes 2-6). THP-1 human monocyte cells were infected with the *Y. pseudotuberculosis* strains. We analyzed YopM translocation into the cells by immunoblotting, and confirmed that the amino-acid substitutions did not impact injection by the T3SS (Fig. S13B, lanes 1-6). Pyrin inflammasome activation was measured in THP-1 cells infected with the *Y. pseudotuberculosis* strains using immunoblotting and ELISA. We monitored the phosphorylation level of pyrin at S242 by immunoblotting and found that the WT YopM proteins maintained total pyrin and S242 phosphorylation, whereas in THP-1 cells infected with *Y. pseudotuberculosis* strains producing the Ala2 variants there was reduced total pyrin and pyrin S242 phosphorylation levels (Fig. 4C, lanes 1-6). As compared to the WT, both YopM^KIM^ Ala2 and YopM^PestA^ Ala2 were deficient for pyrin inflammasome inhibition as measured by release of IL-1β from the THP-1 cells (Fig. 4D). Results of colony-forming unit (CFU) assays performed on bacteria before and after infection indicated that the THP-1 cells were not bactericidal (Fig. S14).

We then tested the contribution of individual side-chain changes in YopM^PestA^ Ala2 on pyrin inflammasome inhibition. We generated *Y. pseudotuberculosis yopJ*^C172A^Δ*yopM* strains producing YopM^PestA^ variants containing alanine substitutions of each of the three residues individually (D159A, D161A or F177A) in Ala2. The strains were tested for YopM^PestA^ secretion as above, and normal levels were confirmed (Fig. S13A, lanes 7-9 vs. lane 5). In the THP-1 infection assay, the YopM^PestA^ variants with single amino-acid changes were translocated at WT levels (Fig. S13B, lanes 7-9 vs lane 5). Mutation of D159 had the strongest impact on the pyrin inhibiting activity of YopM^PestA^, as pyrin dephosphorylation and IL-1β release in THP-1 cells infected with this variant was similar to Ala2 or the empty vector control (Fig. 4C, lane 7, and 4D). The D161 variant had an intermediate phenotype, while mutation of F177 did not have a major impact on YopM^PestA^ pyrin inhibiting activity (Fig. 4C, lanes 8 and 9, and 4D). These data indicate that D159 is essential and D161 is important for YopM^PestA^ to inhibit the human pyrin inflammasome.

### Ala2 variants of YopM^KIM^ and YopM^PestA^ Inhibit the Mouse Pyrin Inflammasome

Results of BACTH and GST pull down assays suggested that YopM^PestA^ and YopM^KIM^ bind with lower affinity to mPYD than hPYD (Fig. 1B, Fig. S3). Based on this result, we hypothesized that binding to mPYD is not required for YopM to target and inhibit mouse pyrin. Therefore, we tested whether YopM Ala2 variants inhibit the pyrin inflammasome during *Y. pseudotuberculosis* infection of murine bone marrow derived macrophages (BMDMs). YopM^KIM^ and YopM^PestA^ Ala2 variants were translocated into infected BMDMs at WT levels (Fig. S13C). YopM^KIM^ and YopM^PestA^ Ala2 variants had only a subtle defect in inhibiting IL-1β release from infected BMDMs, a phenotype that was not statistically significant when compared to WT controls (Fig. S15). Fig. 3D and E and Fig. S9C and D show that in hPYD, R39, R42 and S43 contribute to most of the interactions with YopM^PestA^. The alignment of hPYD with mPYD reveals that while R42 is conserved, position R39 is K39 (conservative change) and S43 is G43 (non-conservative change) in mPYD (Fig. S16A). These and other differences in the binding interface residues between hPYD and mPYD (Fig. S16A, B) are likely responsible for the weak interaction of the latter with YopM.

### The hPYD R42W variant is defective for YopM binding

Amino-acid changes R39G or R42W in hPYD have been implicated in causing FMF, although neither mutation has been validated to be disease causing (50). FMF is an autoinflammatory disease resulting from gain of function mutations in the pyrin gene *MEFV* (25). People with FMF experience recurring episodes of fever with inflammation of serosal membranes in the abdomen, lungs and heart (25). Results of population genetic analysis are consistent with the hypothesis that historic plague epidemics in the Mediterranean selected for FMF gain of function codon changes in the B30.2 domain (39). Since hPYD R42 makes substantial contacts with YopM^PestA^ (Fig. 3D and E) we tested the impact of the R42W codon change in the BACTH. As shown in Fig. 5A, hPYD R42W showed significantly reduced interaction with YopM^PestA^ and YopM^KIM^. Immunoblotting of soluble and insoluble fractions of the *E. coli* strains used in the BACTH was performed to confirm production of hPYD R42W. The hPYD R42W-T18 fusion protein was present in the insoluble samples at comparable amounts to hPYD-T18 (Fig. 5B, lower panel). Lower amounts of hPYD R42W-T18 was seen in the soluble samples as compared to hPYD-T18, consistent with reduced solubility of the PYD domain in *E. coli* in the absence of YopM binding (Fig. S3). Thus, results with the non-conservative R42W change confirms that YopM makes a key interaction with this position, which has been implicated in FMF and positive and negative regulation of the hPYD (51).

**Fig. 5.**
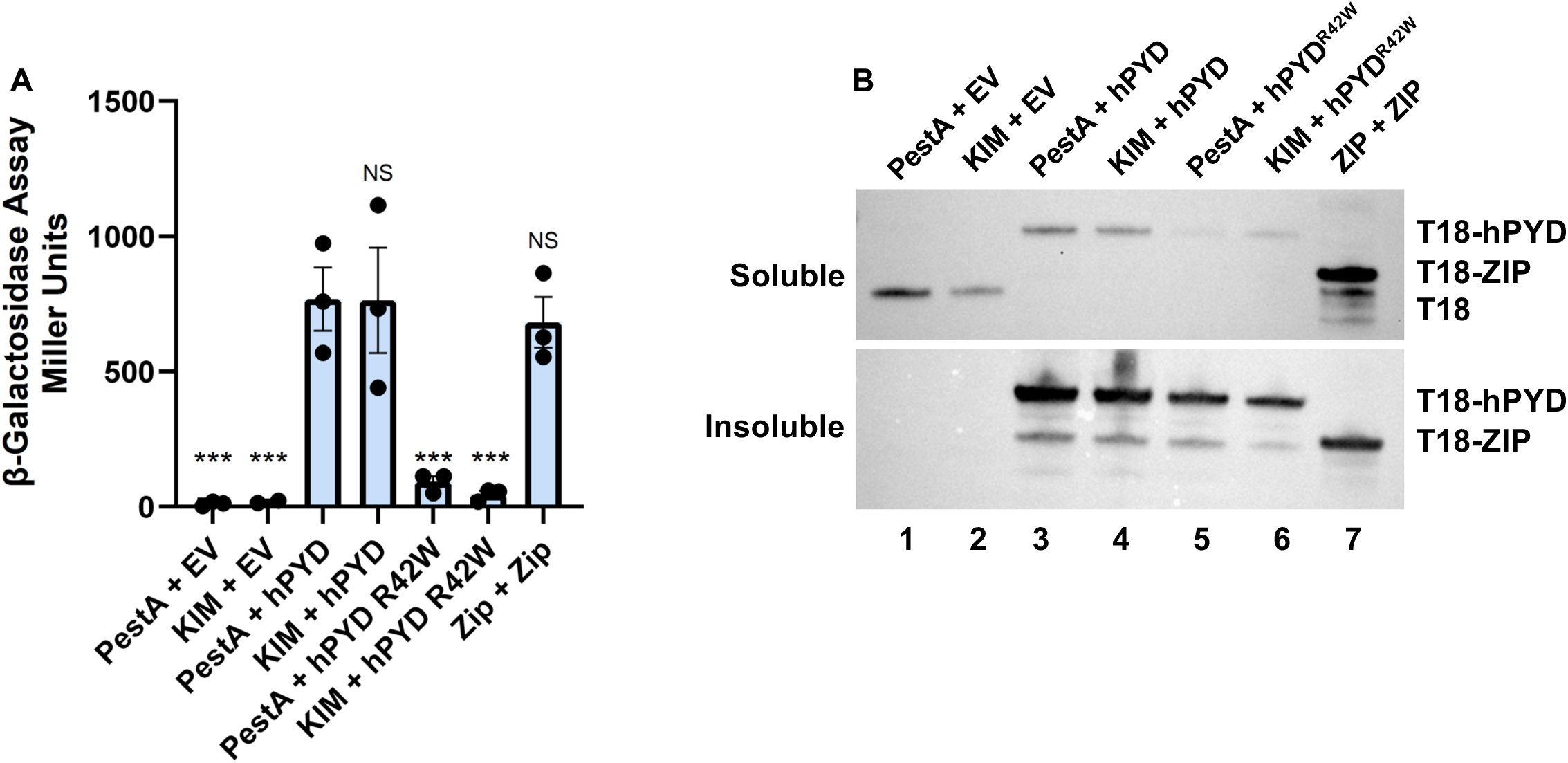
Analysis of hPYD variants by BACTH assay. **A**) The indicated YopM proteins were N-terminally tagged with T25, and hPYDs were C-terminally tagged with T18. Plasmids encoding the fused partners, ZIP positive controls or empty vector (EV) controls were co-transformed into *E. coli* to test for BACTH interactions as in Fig.1. Significance determined by one-way ANOVA followed by a Dunnett’s multiple comparisons test in comparison to PestA + hPYD. ***p<0.001; ns, not significant. **B**) The *E. coli* strains analyzed in (**A**) were lysed and separated into soluble and insoluble fractions. Immunoblot analysis of the fractions was performed with anti-T18 antibody. Results are representative of three independent experiments.

## Discussion

Here, we aimed to investigate the molecular mechanism of the YopM-pyrin interaction. Previous work showed that *Y. pestis* YopM binds human pyrin on the N-terminus that includes both the PYD and linker region (39). We found that *Y. pestis* YopM specifically binds the hPYD, and that the previously identified interaction with part of the linker (39) may have been affected by non-specific binding to GST. *Y. pestis* YopM binding to the hPYD was demonstrated *in vitro* using BACTH, GST pulldowns, and purified proteins. We also determined two co-crystal structures of YopM^PestA^ bound to hPYD and identified residues on both proteins that participate in binding. We show that *Y. pestis* YopM variants which lose binding to hPYD *in vitro* can no longer inhibit the pyrin inflammasome in human monocytes infected with *Yersinia*. Taken together, these data support a model in which *Y. pestis* YopM hijacks host kinases RSK and PRK, binds to the hPYD as a targeting site and promotes pyrin inactivation by phosphorylation of the linker (Fig. S1C).

The YopM^PestA^-hPYD complex is an additional notable structure of a virulence factor targeting a member of the death domain fold (DDF) family of proteins (52). Enteropathogenic *Escherichia coli* encodes T3SS effector NleB, which attaches N-acetylglucosamine to the death domain of the Fas-associated death domain (FADD) or Tumor necrosis factor receptor type 1-associated death domain (TRADD), preventing interaction with death receptors and assembly of death receptor complexes (52).

The DDF family is comprised of four sub-groups; death domain (DD), death effector domain (DED), caspase activation and recruitment domain (CARD) and pyrin domain (PYD) (53). Members of this family play roles in cytosolic innate immune and cell death pathways (53). All DDF proteins participate in homotypic protein-protein interactions resulting in the assembly of higher-order oligomers (53). DDF proteins have a conserved Greek-key motif comprising of six α-helices with variations in the helices defining the sub-families (53). PYDs stand out in this superfamily because they have the shortest helix 3, which is also preceded by a long unstructured α2-α3 loop (51, 54). Helix 3 and the unstructured α2-α3 loop have been hypothesized to play a role in protein-protein interactions of PYDs (51, 54). Intriguingly, R42, which is located in α3, is predicted to be a key residue involved in YopM binding, alongside R39 (Fig. 3DE), which is located on the α2-α3 loop. R42 was proposed to act as a conformational switch due to the unstructured nature of α3 and the proximal α2-α3 loop (54).

The complex structure presented here reveals a strong charge complementarity between YopM^PestA^* and hPYD*. This is not surprising, given that the concave surface of YopM^PestA^* is negatively charged (Fig. 3B) and the hPYD* interacts via a positively charged surface. Three negatively charged residues on YopM^PestA^* (D159, D161 and D203) were predicted from the complex to contribute to hPYD* binding (Fig. S9B). YopM^PestA^* D159 and D161 are located in LRR 5, while D203 is in LRR 7 and was not subjected to detailed analysis in this study. Substitution of YopM^PestA^ D159 had a strong negative impact on hPYD binding, and this could be because it forms hydrogen bonds with R42 and S43 and is also facilitating hydrogen bonds with one of the structural waters observed at the YopM^PestA^*-hPYD* interface (Fig. 3D). Surprisingly, YopM^PestA^* D161 makes extensive interactions with hPYD*, forming two salt bridges with hPYD* R39 and R42 (Fig 3E), yet the impact of its mutation was intermediate.

This might be because the same hPYD* residues make more interactions with other YopM* residues, hence only dampening rather than abolishing the interface. Substitution of YopM F177 had minimal impact on the inhibitory activity of YopM and is unsurprising given the interaction is restricted to more superficial van der Waals contacts with hPYD* (Fig 3F). Three residues on YopM^PestA^* (N120, Y135 and N140) are also predicted to contribute extensive van der Waals interactions and hydrogen bonds yet simultaneously mutating them (Ala1) had no impact on hPYD binding in BACTH. These results could indicate that the overall impact of losing these interactions in the Ala1 variant is minimal, and that YopM relies more on the centrally located acidic residues targeted in Ala2.

Our results indicate that most of the concave surface of the YopM LRR domain is sterically occluded when bound to hPYD. YopM LRRs 6-15 have been implicated in binding to PRK (40). If PRK also binds to the concave surface of the YopM LRR domain, binding to the PRK kinase domain and hPYD would be mutually exclusive. As shown in Fig. S1C, PRK may need to be released to allow YopM to target the hPYD and use hijacked RSK to phosphorylate the human pyrin linker. Additional studies are needed to understand if binding of PRK and hPYD to YopM are mutually exclusive.

PYDs are known to interact through electrostatic interactions (51, 55). Critical to assembly of the pyrin inflammasome is an interaction between the pyrin and ASC PYDs (51) which then allows a recruitment of pro-caspase-1 (56). The two PYDs interact using complementary positive and negatively charged sites (51). Additionally, because these PYDs have multiple charged surfaces, they can simultaneously self-associate and participate in inter-molecular interactions, leading to the assembly of a multi-protein complex (51). Vajjhala *et al*. identified three and two binding sites on pyrin and ASC PYDs, respectively (51). R42 of pyrin PYD and R41 of ASC PYD, which are in α3, and in Site 2 for each protein, were identified to be important for binding to each other (51). The finding that YopM specifically binds to the positively charged patch of hPYD containing R42 and representing Site 2 is intriguing, given that it overlaps with an ASC binding site and the B-box binding site (51). This might indicate that YopM evolved to target a specific binding region on hPYD essential for its function. Additionally, YopM may have to compete with ASC and the B-box to bind hPYD. Future studies that define hPYD interactions with the B-box, ASC and YopM will be informative in testing this hypothesis.

The hPYD R39G and R42W variants have been implicated in FMF, and we found that the latter codon change significantly reduced YopM binding (Fig. 5). The impact of the R42W substitution on hPYD structure and function has been studied, and it was found to increase the stability of the short helix α3 (51). While counterintuitive, this increased structural stability might be deleterious to the function of hPYD, due to the importance of flexibility in this region. The R42W substitution appeared to cause a greater reduction in binding of the B-box to hPYD as compared to ASC (51). The R42W substitution may reduce negative regulation by the B-box (31) which in turn could increase activation of pyrin (51). Recently, Bronnec et al. carried out a high-throughput analysis of *MEFV* variants and reported that neither R39G nor R42W could be validated as pathogenic (57). If the the R42W substitution was selected to reduce YopM binding to hPYD, it is unlikely to have gain of function activity as an added benefit. Most pathogenic FMF codon changes are located on the C-terminal B30.2 domain (25, 50). FMF amino-acid acid changes in the B30.2 domain may overcome YopM inhibition by decreasing the activation threshold of the pyrin inflammasome (58). A second class of gain of function *MEFV* mutations are those that prevent phosphorylation of the linker, such as S242R, resulting in the disease pyrin-associated autoinflammation with neutrophilic dermatosis (59). It is possible that by targeting the key R42-containing site of positive regulation in hPYD YopM forced *MEFV* to evolve gain of function mutations elsewhere, i.e. in the linker or B30.2 domain to counteract plague infection. Additional experiments are needed to understand how the R42W and R39G substitutions in hPYD impact YopM and B-box binding, pyrin activation and *Yersinia* pathogenesis.

The finding that *Y. pestis* YopM does not strongly bind to mPYD could mean that this domain has evolved reduced affinity for recognition. This is possible, given that rodents have likely been hosts for *Y. pestis* longer than humans. Human and mouse PYDs have a sequence identity of 76% and amino-acid differences between the two could indicate sites of positive selection on mPYD. hPYD R39, R42, and S43 have the most interactions with YopM, and two of these sites have conservative (R39 to K39) or non-conservative (S43 to G43) changes in mPYD. These substitutions could be sufficient to disrupt YopM binding. The R47M change in mPYD could also interfere with the band of positively charged surface residues seen in hPYD. Given that there are only three positively charged residues in the YopM binding region (R39, R42 and R47), the loss of one of them might have a disproportionate impact, especially because the PYD is small. Additionally, because the three substituted residues are located on α3, they might cause structural changes that decrease recognition by YopM. These three residues might be the hotspots of mPYD evolution, but a more detailed analysis of positive selection in mouse pyrin would be informative in testing this hypothesis. There is evidence that a small number of residue substitutions can have a significant impact on the interactions of host and pathogen proteins (60). These residues function as hotspots in the evolutionary host-pathogen arms race (60).

Despite the lack of stand-alone mPYD binding, we found that *Y. pestis* YopM Ala2 variants retain the ability to inhibit the pyrin inflammasome in infected mouse macrophages. These results suggest that YopM targets and inhibits murine pyrin independent of the PYD. PRK was shown to be important for YopM to inhibit the pyrin inflammasome in mouse macrophages infected with *Y. pseudotuberculosis* (15). PRK uses its kinase domain to target and phosphorylate pyrin (33, 39). One possibility is that YopM uses hijacked PRK to target and phosphorylate murine pyrin (Fig. S1C). Further studies that focus on how YopM targets and inhibits mouse pyrin are needed to better understand differences in this key mechanism of pathogenesis in murine vs human cells.

## ACKNOWLEDGMENTS AND FUNDING

Supported by NIH grants R01-AI099222 (to JB), P20-GM113132 (bioMT cores; to DM), and T32-AI007519 (training grant stipend to ARS).

This research used 17-ID-1 of the National Synchrotron Light Source II, a U.S. Department of Energy (DOE) Office of Science User Facility operated for the DOE Office of Science by Brookhaven National Laboratory under Contract No. DE-SC0012704. The Center for BioMolecular Structure (CBMS) is primarily supported by the National Institutes of Health, National Institute of General Medical Sciences (NIGMS) through a Center Core P30 Grant (P30-GM133893), and by the DOE Office of Biological and Environmental Research (KP1605010).

## MATERIALS AND METHODS

### Protein Expression Vectors

The vectors and bacterial host strains used are provided in Table 2. In brief, YopM^PestA^ constructs were cloned as N-terminal hexahistidine-GST (6xHisGST) fusions. In addition to the full-length coding sequence (6xHisGST-YopM^PestA^) that was cloned into a modified pGEX_KT vector, a truncated construct lacking the disordered N– and C-terminal extensions (residues 34-344, YopM^ι1Nι1C^) was cloned into a pET28-derived expression vector. These extensions were excluded from the final construct to mitigate potential interference during crystallization trials. The CDS of the human PYD (residues 1-90, herein referred to as hPYD) was cloned as either an N-terminal hexahistidine fusion in a pDuet vector, or as a C-terminal octahistidine fusion in a pET28a vector. All inserts were first PCR amplified then inserted into their respective vectors by either restriction cloning using BamHI and EcoRI sites (pGEX_KT YopM^PestA^), EcoRI and HindIII (pACYCDuet_hPYD) or using the NEB Gibson Assembly kit according to the manufacturer’s recommendations.

### Bacterial Two Hybrid Assay

PYDs and pyrin linker constructs were expressed from pUT18 with C-terminal fusions of T18, and YopM isoforms and variants were expressed from pKT25 with N-terminal fusions to T25. A Quickchange kit (Agilent) was used to introduce codon changes into the BACTH vectors, followed by sequence validation. The BACTH assay was performed as previously described (61, 62). Briefly, electrocompetent BTH101 *E. coli* cells were co-transformed with each vector or empty vector controls and recovered in 250 µl SOC media at 37°C. Cells were plated on an Ampicillin (Amp) or Carbencillin (Carb) + Kanamycin (Kan) plate for 2 days at 30°C and resulting colonies streaked onto Amp/Carb + Kan + isopropyl β-D-1-thiogalactopyranoside (IPTG) plates with and without X-Gal. After 1 day at 30°C, the cells on plates without X-Gal were scraped off and resuspended in Phosphate Buffered Saline (PBS; 140 mM NaCl, 2.68 mM KCl, 8.1 mM Na_2_HPO_4_, 1.47 mM KH_2_PO_4_) and normalized to 0.D_600_ ∼ 0.5. The cells were then incubated at 30°C for 5 min for lysis with 25 µl of 0.1% SDS, 50 µl chloroform in Z buffer (60 mM Na_2_HPO_4_, 40 mM NaH_2_PO_4_,10 mM KCl, 1 mM MgSO_4_) with 50 mM β-mercaptoethanol (β-ME). 200 µl of 4 mg/mL *ortho*-nitrophenyl-β-D-galactopyranoside (ONPG) in Z buffer was added to the cells and the reaction performed for 5 minutes at 30°C. A 500 µl solution of 1 M sodium bicarbonate was added to stop the reaction and the absorbance was measured at 420 and 550 nm. The β-galactosidase activity was then calculated using the following formula as described by Miller:

Miller Units =1000*[(OD_420_ – 1.75*OD_550_)/ (time in minutes*OD_600_*volume of scraped cultures)]

For immunoblotting bacteria recovered from plates as above were lysed with Bugbuster Protein Extraction Reagent (Sigma-Aldrich) and separated into soluble and insoluble fractions by centrifugation. Immunoblotting of soluble and insoluble samples was performed as described below using Cya A (3D1) mouse monoclonal (1:10,000, Santa Cruz Biotechnology).

### GST Pulldown Assay

GST-YopM and His-tagged murine or human PYDs were co-expressed in *E. coli* BL21 cells. The bacterial lysates were prepared with BugBuster^®^ Protein Extraction Reagent (Millipore) in PBS and supplemented with lysonase (Millipore) and Roche complete mini protease inhibitor. Lysates were then incubated with GST resin (GST.Bind, Millipore) overnight. The resin was then washed and proteins eluted using 1X sample buffer and analyzed on SDS-PAGE gels followed by blue staining or α-His immunoblotting. Immunoblotting procedures are described below.

### Protein Expression and Purification

All recombinant proteins used in this manuscript were expressed in BL21(DE3) cells. Proteins were either co-expressed and co-purified for crystallization of the YopM^PestA^*-hPYD* structure or individually prepared for the YopM^ΔNΔC^-hPYD complex. In all preparations, cells were grown in Terrific Broth (TB) at 37°C for 2-3 hours until an OD_600_ of ∼ 1 was achieved. Cells were induced with 0.1 mM IPTG and protein expression was allowed to proceed at 18°C for 16-18 h. For YopM, induced cells were allowed to express protein at 37°C for 4 h. Cells were harvested by centrifugation at 4,200 x *g* at 4°C for 15 min. Cells were resuspended in lysis buffer (150 mM NaCl, 1 mM Tris(2-carboxyethyl)phosphine hydrochloride [TCEP], and 20 mM Tris, pH 7.4) supplemented with Roche complete mini EDTA (ethylenediaminetetraacetic acid) protease inhibitor and Lysonase Bioprocessing Reagent from Millipore Sigma and lysed using an LM10 microfluidizer. The insoluble fraction was separated by centrifugation using a Ti 45 Rotor in a Beckman Ultima L-70 ultracentrifuge at 40,000 rpm for 1 h at 4°C. For both the YopM^PestA^ only construct and YopM^PestA^ co-expressed with hPYD, recombinant protein was captured on GST resin and eluted by on-column proteolysis using the Tobacco Etch Virus (TEV) protease in buffer composed of 150 mM NaCl, 1 mM TCEP, and 20 mM Tris, pH 7.4. For YopM co-purified with hPYD, the hexahistidine tag on hPYD was cleaved using the HRV-3C protease during dialysis against buffer containing 150 mM NaCl, 1 mM TCEP, and 20 mM Tris, pH 7.4; cleavage and dialysis were allowed to proceed overnight at 4°C. In both cases, recombinant protein was further purified by size exclusion chromatography (SEC) using a HiLoad Superdex 200 26/600 PG column and fractions containing purified protein, as assessed by reducing SDS-PAGE, were pooled and concentrated.

For the YopM^ΔNΔC^-hPYD complex structure, proteins were individually expressed and purified. The bacterial pellets were expressed, lysed, and clarified as described above except using a 20 mM 4-(2-hydroxyethyl)-1-piperazineethanesulfonic acid (HEPES), pH 7.4 buffer system in place of Tris. Clarified lysates were applied to a 5 mL His-TRAP column then washed with buffer composed of 500 mM NaCl and 20 mM Tris, pH 8.0 supplemented with imidazole concentrations ranging from 10-50 mM. Proteins were eluted by gradient elution ranging from 10-250 mM imidazole. For the hPYD preparation, fractions containing protein were pooled, concentrated and run on a HiLoad Superdex 200 26/600 PG column in buffer composed of 150 mM NaCl, 1mM TCEP, and 20 mM HEPES, pH 7.4. For YopM^ΔNΔC^, fractions containing recombinant protein were pooled and TEV cleavage reactions were set up during simultaneous dialysis against buffer composed of 150 mM NaCl, 1 mM TCEP, and 20 mM HEPES, pH 7.4 at 4°C, for 16-18 h. Cleaved protein was further purified by cation exchange using SP Sepharose and IMAC using Ni-NTA resin. Finally, samples were purified by SEC using a HiLoad Superdex 200 26/600 PG column.

### Analytical SEC and SEC-MALS

Analytical SEC was performed on a Superdex 200 10/300 column attached to an ÄKTA pure™ protein purification system (Cytiva). Samples were analyzed in 150 mM NaCl, 1 mM TCEP, and 20 mM Tris, pH 7.4 for GST resin purified YopM^PestA^ and YopM^PestA^-hPYD. YopM^ΔNΔC^ and YopM^ΔNΔC^-hPYD complex were analyzed in 150 mM NaCl, 1 mM TCEP, and 20 mM HEPES, pH 7.4. Analytical SEC was performed at 0.75 mL/min. The column was calibrated using the 29,000-700,000 Da gel filtration markers kit from Millipore Sigma (MWG1000F) containing the following molecular mass standards: Blue Dextran (2,000 kDa), Apoferritin (440 kDa), β-Amylase (200 kDa), Alcohol Dehydrogenase (150 kDa), Albumin (66 kDa), Carbonic Anhydrase (29 kDa).

For SEC-MALS, an ÄKTA pure™ instrument (Cytiva) coupled to an Optilab refractive index detector (Wyatt) and light-scattering detector (Wyatt MiniDawn Tristar) were used. SEC was performed using a Superose 10/300 Column at 0.5 mL/min and the laser wavelength was set to 690 nm. The molar mass was determined using the ASTRA software package (Wyatt).

### Velocity Ultracentrifugation

Analytical ultracentrifugation was conducted using a Beckman Proteomelab XL-A and an AN-60 rotor. For sedimentation velocity analytical ultracentrifugation, YopM^PestA^ in 150 mM NaCl, 1 mM TCEP, with either 20 mM Tris, pH 7.4 or 20 mM MES pH 5.0 was centrifuged at 35,000 rpm with monitoring at 280 nm. Data were analyzed using Sedfit (63) to determine sedimentation coefficient, frictional ratio, and apparent mass, as described elsewhere (64). The sedimentation coefficient reported is that of the major peak (at least 80% of the total analyzed mass) at OD_280_ for each sample.

### Crystallization

The YopM^PestA^*-hPYD* complex was crystallized using the co-purified YopM-hPYD. The complex was crystallized by vapor diffusion in 4 µL sitting drops composed of well solution (12% [*w/v*] PEG 4K, 40 mM MES, pH 5.0) and co-purified YopM^PestA^-hPYD at a concentration of 6.5 mg/mL as determined by the Bradford assay; drops were set at a 1:1 ratio. Prior to harvesting, crystals were cryoprotected in well solution supplemented with 20% (*v/v*) glycerol then flash cooled in liquid nitrogen.

For co-crystallization of the YopM^ΔNΔC^-hPYD complex, the purified YopM^ΔNΔC^ was first combined with purified hPYD in a 1:4 molar ratio at a final total protein concentration of 1.2 mg/mL. Suitable crystals were obtained by vapor diffusion in 4 µL hanging drops composed of well solution (19% [*w/v*] PEG 3350, 300 mM NaSCN, and 100 mM Bis-Tris Propane, pH 6.5) and protein set at a 1:1 drop ratio. Crystals were harvested and flash cooled as described above.

### Structure Determination and Refinement

For both co-crystal structures, data were collected at the National Synchrotron Light Source II on the AMX beamlines (17-ID-1) using a Dectris 9M pixel detector. For the full-length co-purified YopM^PestA*^-hPYD* complex, 180° of rotation X-ray diffraction data were collected using 0.1° per frame. For the YopM^ΔNΔC^-hPYD complex, 120° of rotation X-ray diffraction data were collected using 0.2° per frame. The diffraction images were reduced using XDS and scaled using XSCALE (65). The R_free_ set was generated from 5% of reflections in thin resolution shells using Phenix reflection file editor. The Matthews coefficient was determined using the CCP4 supported program MATTHEWS_COEF (66). In both cases, initial phases were determined by molecular replacement using PHASER-MR and iterative automated and manual refinements were carried out using phenix.refine and Coot, respectively. Specific details pertaining to the phasing and refinement of each structure follows.

For the YopM^PestA^*-hPYD* complex, initial phase estimates were obtained using the crystal structure of NLRP9 PYD (pdb_00006z2g) as a search model. For the 13-LRR YopM^PestA^ isoform used in these studies, a composite model was initially created using the crystal structure of a related 15-LRR *Y. pestis* YopM isoform (pdb_00001g9u) in PyMol (67) to omit additional leucine rich repeats in the central LRR domain. Following molecular replacement, negative density peaks were observed in the m*Fo* – d*Fc* map along the entirety of the C-terminal portion of YopM. After omission of the region spanning residues 248-344 followed by iterative automated and manual refinements using phenix.refine and Coot, respectively, it was determined that a YopM^PestA^ truncation was captured in the crystal lattice. We posit a limited proteolysis event occurred in the mother liquor preceding nucleation that resulted in a shortened YopM^PestA^ in the co-purified YopM^PestA^-hPYD construct represented in the deposited coordinates. This conclusion is supported by the inability of the missing YopM^PestA^ sequence to be included in either molecule of the asymmetric unit without incurring severe steric overlap with molecules related by both crystallographic and non-crystallographic symmetry. A region spanning residues 79-90 and encompassing the last α-helix of hPYD is also absent in this crystal structure and is presumed to be another target of errant proteolytic activity. The stable complex fragments resolved in this co-crystal structure are denoted as YopM^PestA^*-hPYD*. The CCP4 supported programs PEAKMAX and WATPEAK (68) were used to identify unmodeled positive density peaks and evaluate them for waters, which were modeled as appropriate following manual inspection. Additionally, a > 4 sigma positive density peak adjacent to the sulfhydryl group of Cys100 of Chain B was observed that exhibited good fitment to a carboxymethyl cysteine. However, LC-MS/MS analysis of crystals rehydrated from neighboring drops with similar well-solution compositions failed to corroborate the observations in the crystallographic data despite evidence for low-abundance carboxymethylation at other cysteines. The positive density peak was therefore left unmodeled. Anisotropic refinement of protein atoms was carried out using one TLS group per chain; subsequent refinements transpired without incident.

For the co-crystal structure of the YopM^ΔNΔC^-hPYD complex, initial phase estimates were obtained using a model composed of the refined YopM^PestA^*-hPYD* structure described above with a graft of the C-terminal portion of the aforementioned *Y. pestis* YopM isoform (pdb_00001g9u, residues 286-396). The graft was created by performing a least-squares superposition of the preceding two LRRs with the corresponding region of the refined YopM^PestA^*-hPYD* complex. Due to the low resolution and map quality, automated refinements were performed using secondary structure restraints. The refined coordinates of the YopM^PestA^*-hPYD* complex were also used to impose reference model restraints during automated refinement but were lifted for the final stages.

### Accession Numbers

The coordinate files and structure factor files for both co-crystal structures solved herein are available at the Protein Data Bank (69) using the following codes: pdb_00009oyu (YopM^PestA^*-hPYD*) and pdb_000010dr (YopM^ΔNΔC^-hPYD). Rotation X-ray diffraction image files were uploaded to the SBGrid Data Bank and are searchable by PDB codes.

### Structural Analyses and Visualization

Structural analyses were performed using PyMol (67) and the CCP4 Suite (68) as part of the SBGrid software environment (70). Structural alignments were performed in PyMol using all C_a_ of the YopM chains for each complex. The Adaptive Poisson-Boltzmann Solver (71) (APBS) plugin was used to calculate electrostatic surface potentials for YopM^PestA^* and hPYD* in PyMol. All structural figures were rendered using PyMol. Residues at the binding interface were identified using the CCP4 supported program CONTACTS (68) with a 5 Å cutoff, and histograms displaying per-residue contacts were created using R (72) with the following packages: tidyverse (73), ggprsim (74), and ggplot2 (75).

### Construction of *Y. pseudotuberculosis* Strains

*Y. pseudotuberculosis* strains expressing different YopM isoforms and variants (see Table 2) were generated by conjugating donor *E. coli* S17 lambda-pir containing the pMMB67EH plasmid plus YopM isoforms/variants with a *Y. pseudotuberculosis* strain lacking the *yopM* gene and harboring the catalytic mutation in YopJ (mJ*ΔyopM*).

### Yop Secretion Assay

*Y. pseudotuberculosis yopJ^C172A^Λ1yopM* strains were inoculated into LB Amp and grown with shaking at 28°C. Overnight cultures of *Yersinia* were diluted 1:40 in LB media containing 20 mM sodium oxalate and 20 mM MgCl_2_ and grown for 2 h at 28°C then shifted to 37°C for 4 h. Supernatants were precipitated using trichloroacetic acid (TCA) overnight. Precipitated proteins were centrifuged at 14,000 rpm for 30 min, and the resulting pellet washed with acetone. The protein pellets were resuspended in 100 µl 1X NuPAGE LDS Sample Buffer and Sample Reducing Agent and 10 µl of the samples were run on 4–12% NuPAGE Bis-Tris SDS-PAGE gels (Invitrogen by ThermoFisher Scientific). The gels were stained using GelCode Blue Stain Reagent (ThermoFisher) or One-Step Blue Protein Gel Stain (Biotium) and imaged on an iBright FL1500 imager (ThermoFisher Scientific).

### THP-1 Cells Tissue Culture and Infections

THP-1 human monocyte cells (ATCC TIB-202) were grown in filter-sterilized RPMI-1640 media with 10% heat-inactivated fetal bovine serum (FBS), 0.05 mM β-ME and 1% pen-strep (10,000 U/mL Penicillin + 10,000 µg/mL streptomycin, Gibco) in 25 cm^2^ flasks. Once the cells reached confluency, they were split into 75 cm^2^ flasks. For infections, 8×10^5^ total cells per well were seeded in 1.5 mL serum-free media in 6-well plates. Overnight cultures of *Yersinia* were diluted 1:40 in LB media containing 20 mM sodium oxalate and 20 mM MgCl_2_ and grown for 1 h at 28°C, then shifted to 37°C for 2 h. Cultures were centrifuged and the bacterial pellets washed in 1X PBS. Resuspended bacteria were then mixed with cells at a multiplicity of infection (MOI) of 30 in serum-free media then plated on 6-well plates. After infection for 4 h at 37°C in 5% CO_2_, the cell contents were collected and centrifuged for 15 min at 14,000 rpm. The supernatants were collected for IL-1β analysis by ELISA (see below). The cell pellet was then washed with 1X Hanks balanced salt solution (HBSS) and lysed for immunoblot analysis as described below. Samples of bacteria inoculated into each well at t=0 h were serially diluted and plated for CFU quantification. Total bacterial-monocyte co-cultures were solubilized in 0.1% NP-40 detergent (t=4 h) followed by serial dilutions and plating to determine CFU recovery post-infection.

### BMDM Cells Tissue Culture and Infections

BMDMs were obtained from femurs and tibia of 8-week-old WT C57BL/6 mice (Jackson Laboratories) and cultured as previously described (76). After differentiation for 7 days, the BMDMs were seeded in 3 mL MGM 10/10 containing Dulbecco’s modified Eagle Medium (DMEM) plus GlutaMAX (Gibco), 10% (*v/v*) heat-inactivated FBS (GE HealthCare), 10% (*v/v*) L929 cell-conditioned medium, 1 mM sodium pyruvate (Gibco), and 10 mM HEPES (Gibco) at a density of 0.8×10^6^ cells/well. The cells were allowed to settle for at least 1 h then primed overnight by addition of 100 ng/mL O26:B6 *E. coli* LPS (Sigma) in 1X PBS. *Yersinia* strains were grown as described above and added to serum-free MGM 10/10. MGM 10/10 with or without bacteria at an MOI of 30 was added to the wells of primed macrophages. Plates were centrifuged for 5 min at 112 x *g* to facilitate *Yersinia* contact with cells and incubated for 90 min at 37°C in 5% CO_2_. Following the infection, cell supernatants were collected for IL-1β analysis by ELISA and cells lysed for immunoblot analysis (see below).

### THP-1 and BMDM Lysates and Immunoblotting

THP-1 and BMDM cells were lysed in mammalian protein extraction reagent (MPER, Thermo Scientific) supplemented with protease inhibitor cocktail and phosphatase inhibitor (Roche). The lysates were spun down to separate out soluble from insoluble fractions. Protein concentration levels were measured using the bicinchoninic acid assay (BCA, Thermo Fisher Scientific). 5-10 µg total soluble protein lysate was run on 4-12% NuPAGE Bis-Tris SDS-PAGE gels (Invitrogen by ThermoFisher Scientific) and transferred to polyvinylidene difluoride membranes (ThermoFisher Scientific) using an iBlot 3 gel transfer device (Life Technologies). Membranes were blocked using 5% (*w/v*) nonfat dairy milk (Lab Scientific bioKEMIX) for 1 h at room temperature before incubation with primary antibody overnight. Primary antibodies used in this study include rabbit anti-mouse phospho-serine 241 monoclonal antibody (1:1,000 dilution, Ab200420; Abcam), rabbit anti-human pyrin monoclonal (1:1,000, Cell Signaling #58525 and #40649), rabbit-anti-mouse/human polyclonal β-actin (1:1,000 # 4967; Cell Signaling), rabbit anti-*Y. pestis* YopM polyclonal (1:10,000 dilution, antibody provided by Susan Straley) and mouse anti-HIS tag monoclonal (1:5,000 dilution, # MMS-156P BioLegend). Horseradish peroxidase-conjugated anti-rabbit or anti-mouse (Jackson Laboratory) were used as secondary antibodies. Protein bands bound by antibodies were visualized using chemiluminescent detection reagent (GE HealthCare) on an iBright FL1500 imager (ThermoFisher Scientific). Band quantification was also performed on the iBright FL1500 imager.

### IL-1β Quantification

Levels of IL-1β in THP-1 and BMDM supernatants were quantified using human (RandD Systems) and mouse (MLB00C, R&D Systems) ELISA kits respectively, following the manufacturer’s instructions.

### Statistical Analysis

GraphPad Prism was used to perform statistical analyses. Significance was determined by one-way ANOVA and Tukey’s *post hoc* tests performed for BACTH and ELISA experiments.

| **Table 2. Vectors Name** | **Bacterial host strain** | **Description or insert** | **Reference** |
| --- | --- | --- | --- |
| pUT18 | BTH101 | pUT18 empty vector allowing fusion to the N-terminus of the cya T18 fragment | Euromedex |
| pKT25 | BTH101 | pKT25 empty vector allowing fusion to the C-terminus of the cya T25 fragment | Euromedex |
| pKT25_YopM<sup>PestA</sup> | BTH101 | Full-length YopM<sup>PestA</sup> CDS | This study |
| pKT25_YopM<sup>PestA</sup> Ala1 | BTH101 | YopM<sup>PestA</sup> Ala1 (N120A, Y135A, and N140A) | This study |
| pKT25_YopM<sup>PestA</sup> Ala2 | BTH101 | YopM<sup>PestA</sup> Ala2 (D159A, D161A, and F177A) | This study |
| pKT25_YopM<sup>PestA</sup> Ala3 | BTH101 | YopM<sup>PestA</sup> Ala3 (Y201A, N221A, and E279A) | This study |
| pKT25_YopM<sup>KIM</sup> | BTH101 | Full-length YopM<sup>KIM</sup> CDS | This study |
| pKT25_YopM<sup>KIM</sup> Ala2 | BTH101 | YopM<sup>KIM</sup> Ala2 (D159A, D161A, and F177A) | This study |
| pUT18_hPYD | BTH101 | Human pyrin aa 1-90 | This study |
| pUT18_hPYD R42W | BTH101 | Human pyrin aa 1-90 R42W | This study |
| pUT18_mPYD | BTH101 | Mouse pyrin aa 1-90 | This study |
| pUT18_hPyrin186-326 | BTH101 | Human pyrin aa 186-326 | This study |
| pKT25_ZIP | BTH101 | Leucine zipper | Euromedex |
| pUT18C_ZIP | BTH101 | Leucine zipper | Euromedex |
| pGEX_2T | BL21 | pGEX_2T empty vector allowing fusion to the C-terminus of the GST tag | Chung et al 2016 |
| pGEX_2T_YopM<sup>KIM</sup> | BL21 | Full-length YopM<sup>KIM</sup> CDS | Chung et al 2016 |
| pACYCDuet_hPYD | BL21 | Human pyrin aa 1-90 | This study |
| pACYCDuet_mPYD | BL21 | Mouse pyrin aa 1-90 | This study |
| pGEX_KT_YopM<sup>PestA</sup> | BL21 | Full-length YopM<sup>PestA</sup> CDS in a modified pGEX_KT vector with a 6XHIS and TEV site downstream of the thrombin site | This study |
| pET_6XHIS_GST_TEV_YopM<sup>ΔNΔC</sup> | BL21 | YopM<sup>PestA</sup> aa 34-344 | This study |
| pET28A_hPYD_8xHIS | BL21 | Human pyrin aa 1-90 in a modified pET28A vector with an 8X HIS tag | This study |
| pMMB67EH_YopM<sup>PestA</sup> WT | *Y. pseudotuberculosis* mJΔyopM | Full-length YopM<sup>PestA</sup> CDS | This study |
| pMMB67EH_YopM<sup>PestA</sup> Ala1 | *Y. pseudotuberculosis* mJΔyopM | YopM<sup>PestA</sup> Ala1 (N120A, Y135A, and N140A) | This study |
| pMMB67EH_YopM<sup>PestA</sup> Ala2 | *Y. pseudotuberculosis* mJΔyopM | YopM<sup>PestA</sup> Ala2 (D159A, D161A, and F177A) | This study |
| pMMB67EH_YopM<sup>PestA</sup> D159A | *Y. pseudotuberculosis* mJΔyopM | YopM<sup>PestA</sup> (D159A) | This study |
| pMMB67EH_YopM<sup>PestA</sup> D161A | *Y. pseudotuberculosis* mJΔyopM | YopM<sup>PestA</sup> (D161A) | This study |
| pMMB67EH_YopM<sup>PestA</sup> F177A | *Y. pseudotuberculosis* mJΔyopM | YopM<sup>PestA</sup> (F177A) | This study |
| pMMB67EH_YopM<sup>PestA</sup> Ala3 | *Y. pseudotuberculosis* mJΔyopM | YopM<sup>PestA</sup> Ala3 (Y201A, N221A, and E279A) | This study |
| pMMB67EH_YopM<sup>KIM</sup> WT | *Y. pseudotuberculosis* mJΔyopM | Full-length YopM<sup>KIM</sup> CDS | This study |
| pMMB67EH_YopM<sup>KIM</sup> Ala2 | *Y. pseudotuberculosis* mJΔyopM | YopM<sup>KIM</sup> Ala2 (D159A, D161A, and F177A) | This study |

2 Bacterial expression vectors used in this study are listed by name, bacterial host strain, description or 3 insert and reference to indicate source.

---

**Fig. S1.**
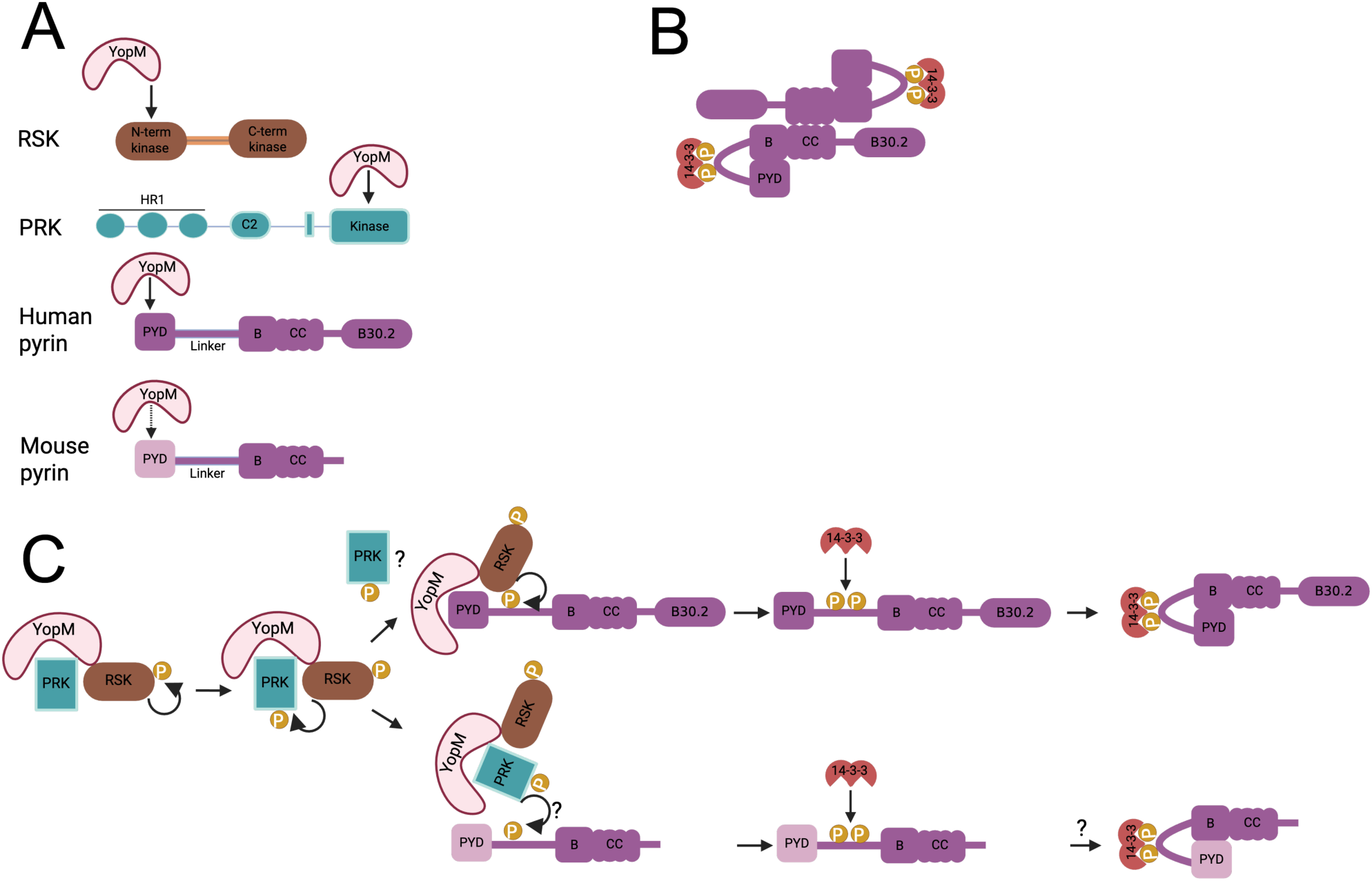
Scheme depicting YopM binding partners and mechanisms of pyrin targeting and inactivation. **A**) Domain arrangements of RSK, PRK, and monomeric human and mouse pyrin and regions targeted for binding by YopM. The C-terminus of YopM binds the N-terminal kinase domain of RSK, and the concave surface of YopM LRR domain binds the kinase domain of PRK and the PYDs of human and mouse pyrin, the latter weakly (dashed arrow). Color shading is used to denote amino acid sequence difference between human and murine PYDs. HR1=homology region 1; C2=C2-like domain; B=B-box; CC=coiled coil. **B**) Anti-parallel dimer of inactive pyrin resulting from coiled-coil interactions. **C**) Model of YopM hijacking and sequentially activating by phosphorylation with RSK and PRK and different targeting mechanisms for human and mouse pyrin. Individual kinase domains of RSK and PRK and monomeric pyrin proteins are shown for simplicity. Phosphorylation of two serine residues in the pyrin linker results in binding of a 14-3-3 dimer and the formation of a hairpin that facilitates sequestration of the PYD by the B-box. To inactivate human pyrin YopM releases PRK prior to PYD binding and uses hijacked RSK to phosphorylate the linker. Role of the released PRK is unknown. To inactivate mouse pyrin YopM uses hijacked PRK to target and phosphorylate the linker. Role of the B-box in negative regulation of mouse pyrin is unknown.

**Fig. S2.**
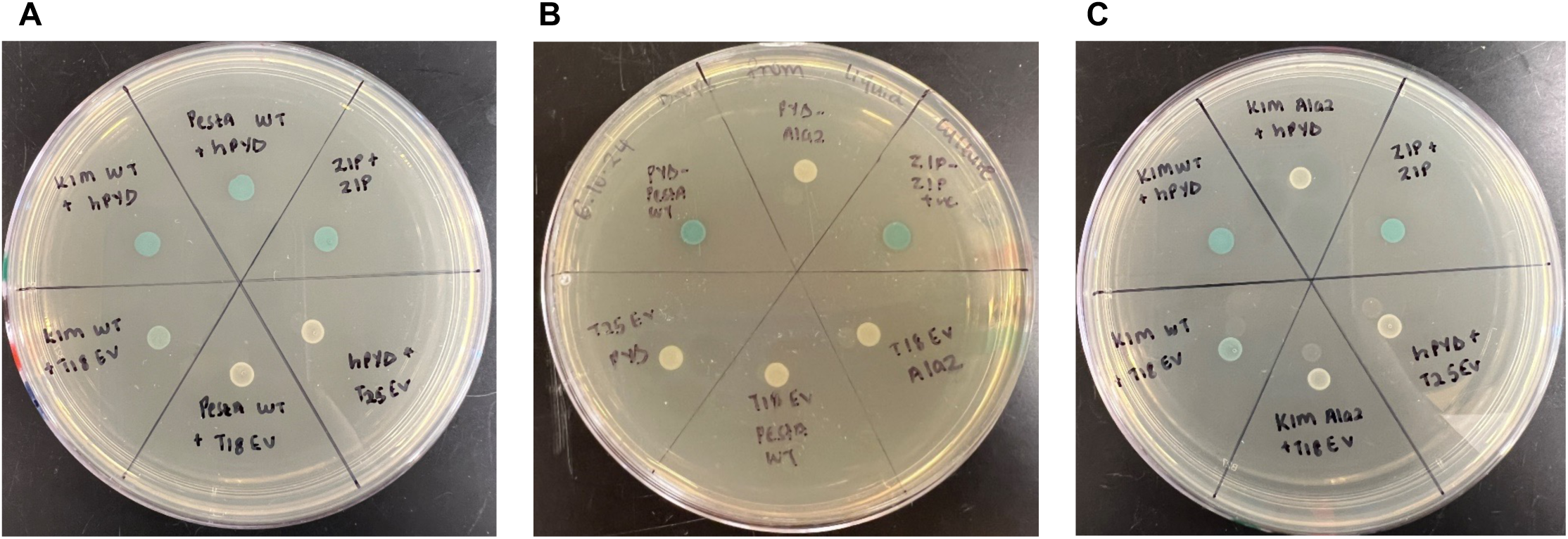
Analysis of YopM-PYD interactions using BACTH. The indicated YopM proteins were N-terminally tagged with T25 and PYDs were C-terminally tagged with T18. Plasmids encoding the fused partners, ZIP positive controls or empty vector (T25 or T18 EV) controls were co-transformed into the *ΔcyaA E. coli* strain BTH101 to test for protein-protein interactions. Suspensions of colonies used for the β-galactosidase assays in Fig.1A (**A**), Fig. 4A (**B**) or Fig.4B (**C**) were dropped on a plate containing X-Gal for qualitative assessment of the protein interactions. Blue color indicates strong interactions and white color depicts weak interactions.

**Fig. S3.**
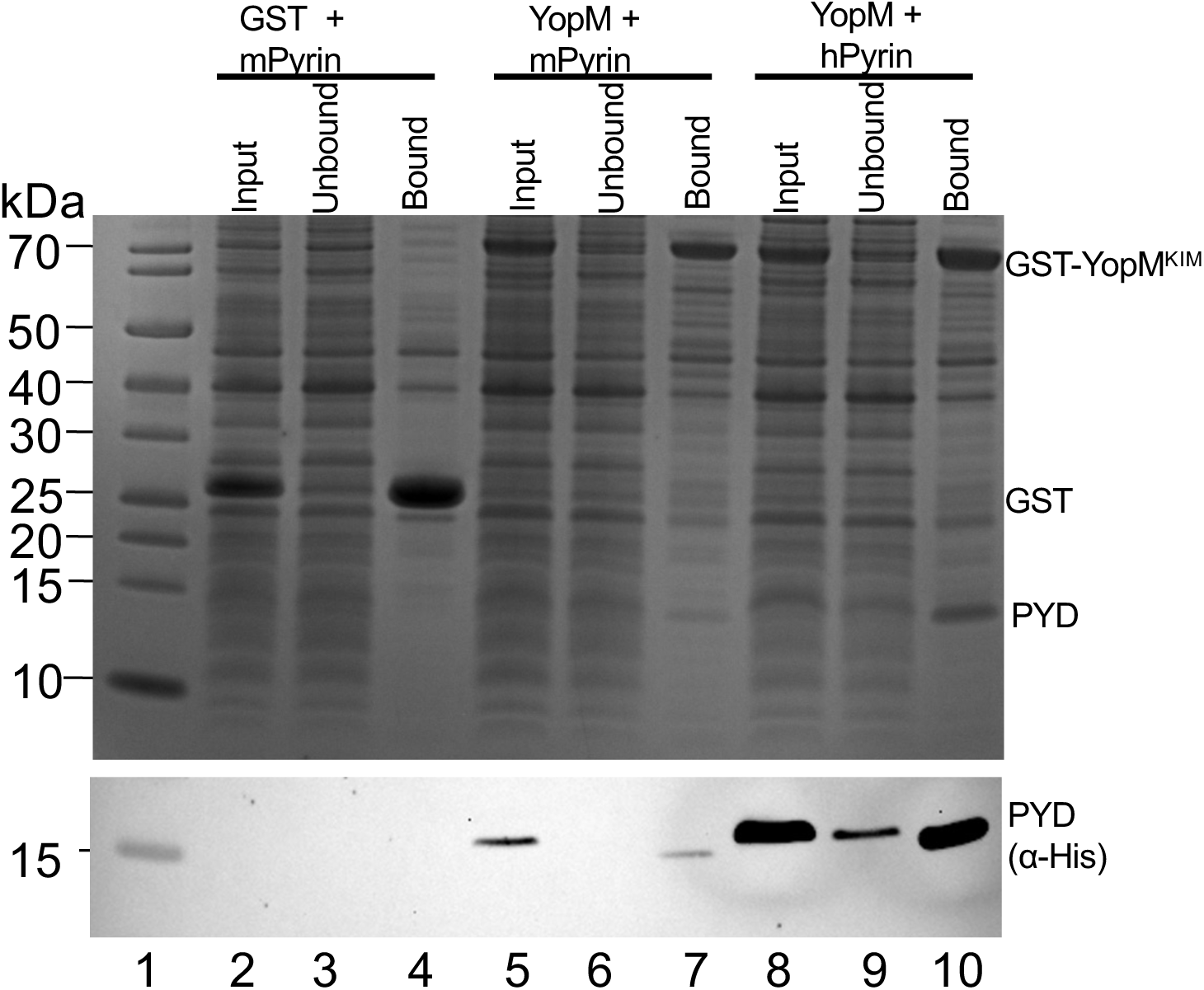
Analysis of YopM-PYD interactions by GST pulldown assay. GST or GST-YopM^KIM^ were co-expressed with His-tagged mPYD or hPYD in *E. coli* and the bacterial lysates were subjected to pull downs using GST resin. Samples of the input, unbound and bound fractions were analyzed by SDS-PAGE and blue protein staining (top) or α-His immunoblotting (bottom). Lane 1 is a M_r_ ladder with sizes labelled on the left. Positions of GST-YopM^KIM^, GST, and PYDs are shown on the right.

**Fig. S4.**
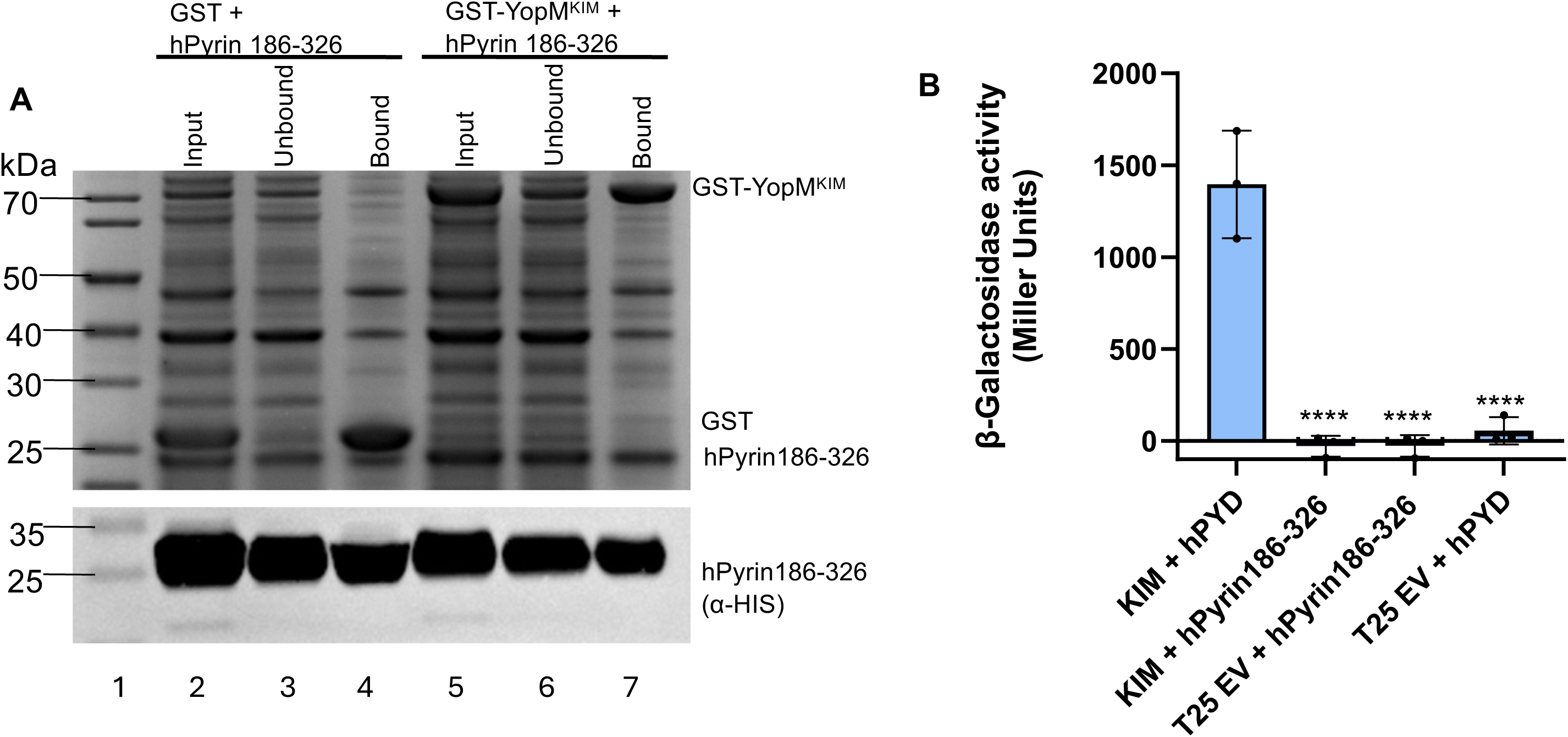
Analysis of YopM-pyrin linker interactions by GST pulldown or BACTH. **A**) GST or GST-YopM^KIM^ were co-expressed with His-tagged hPyrin186-326 in *E. coli* and the bacterial lysates were subjected to pull downs as in Fig. S3. **B**) BACTH analysis of YopM-hPyrin186-326 interactions. YopM^KIM^ was N-terminally tagged with T25 and hPYD or hPyrin186-326 were C-terminally tagged with T18. Plasmids with the fused partners or empty vector (EV) controls were co-transformed into *E. coli* to test for protein-protein interactions as in Fig.1. Results are pooled from at least three independent experiments. Significance determined by one-way ANOVA and Tukey’s post hoc test comparing to KIM + hPYD; ****p<0.0001.

**Fig. S5.**
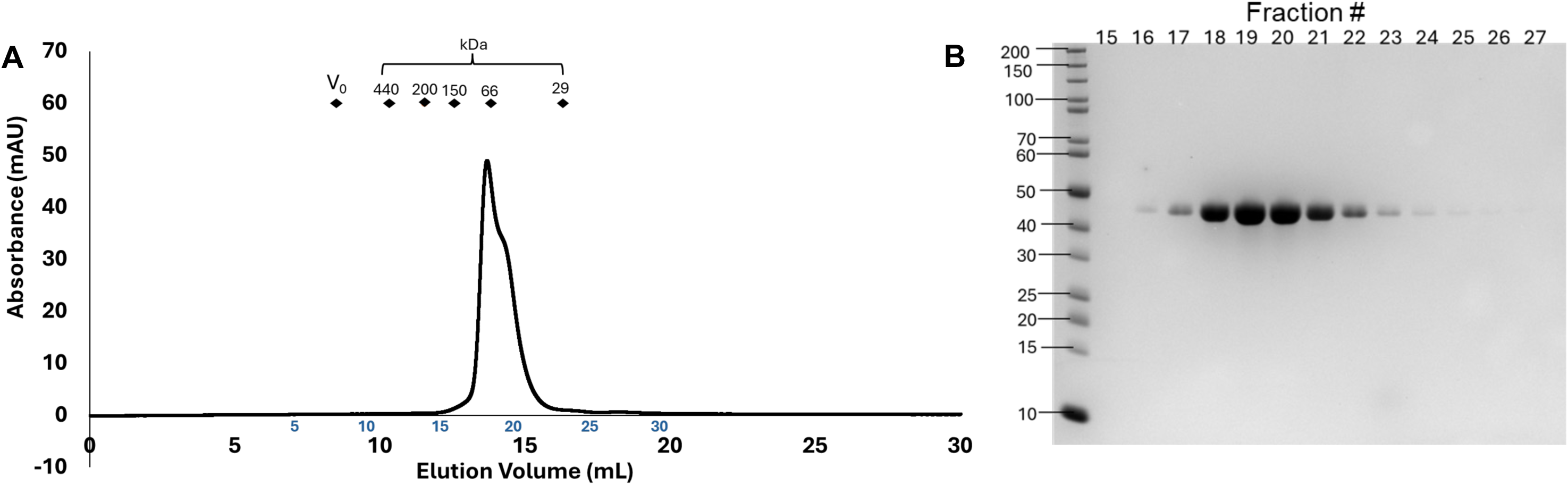
Characterization of YopM^PestA^ by SEC. YopM^PestA^ was analyzed by SEC on a Superdex 200 10/300 column at 0.75 mL/min. Samples were analyzed in buffer containing 20 mM HEPES, 150 mM NaCl and 1 mM TCEP, pH 7.4. **A**) SEC chromatogram with fraction numbers shown in blue on the chromatograms above the elution volumes. Diamonds indicate the column void volume (V_0_) and peak elution volumes of molecular mass standards (440 kDa = apoferritin, 200 kDa = β-amylase, 150 kDa = alcohol dehydrogenase, 66 kDa = albumin, 29 kDa = carbonic anhydrase). **B**) Fractions of the peak eluates were analyzed by SDS-PAGE and blue protein staining.

**Fig. S6.**
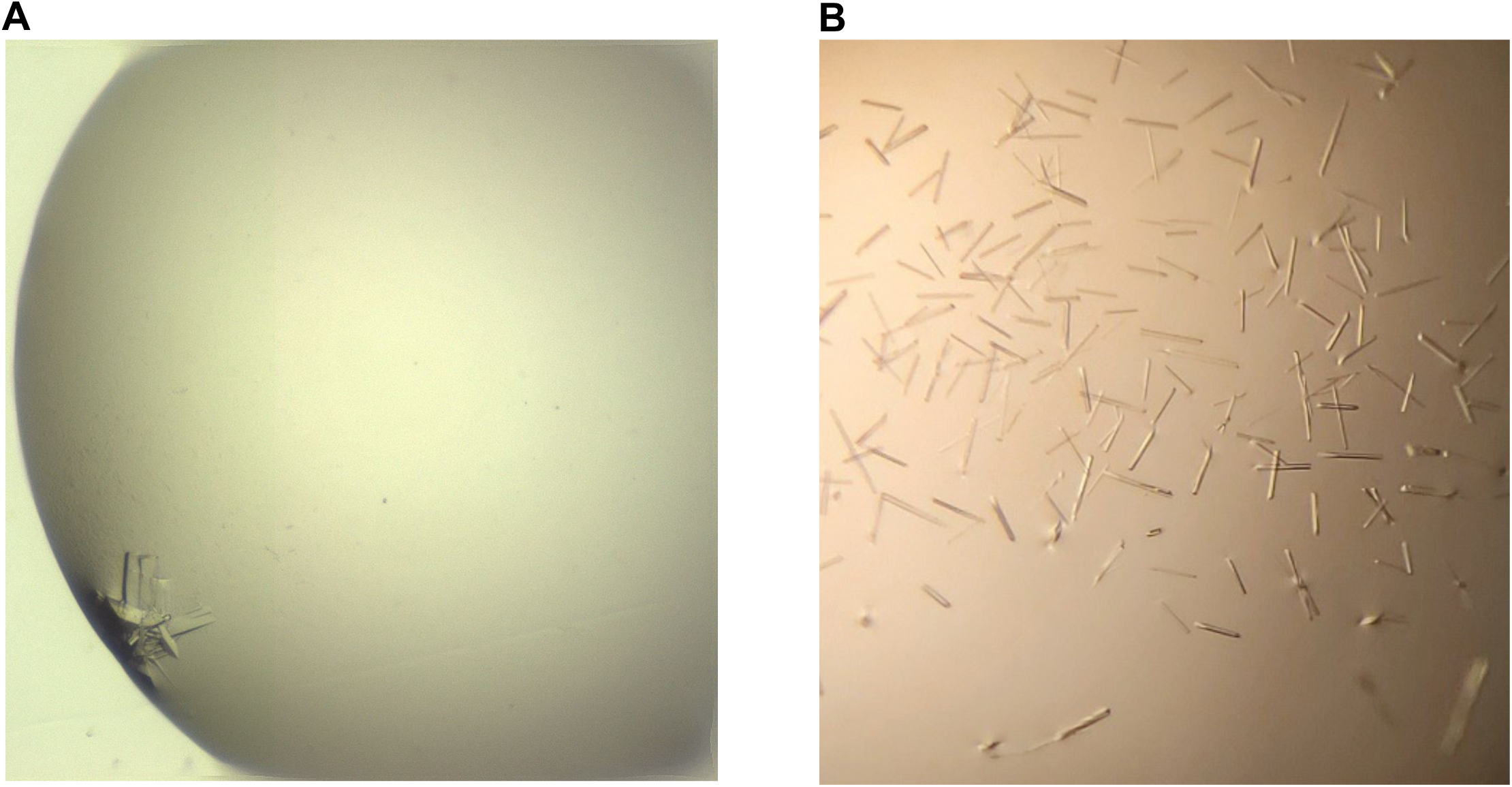
Representative images of YopM-hPYD complex crystals. **A)** YopM^PestA^ + hPYD crystals in 12% PEG 4K,0. MES, pH5.0 growing in sitting drops**. B**) YopM^ΔNΔC^ + hPYD crystals in 19% PEG 3350, 0.3 M NaSCN, 0.1 M Bis-Tris propane, pH 6.5 growing in hanging drops.

**Fig. S7.**
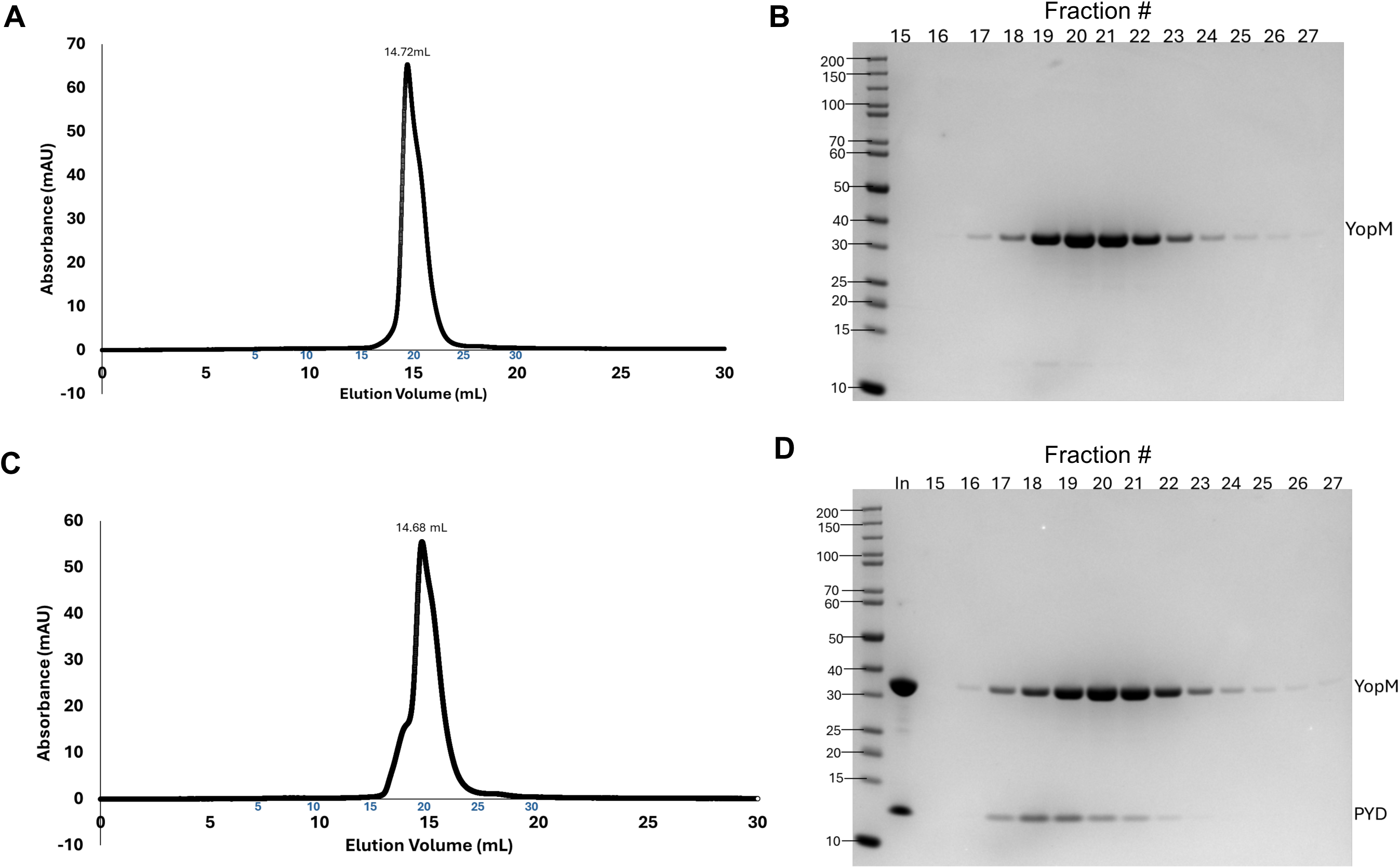
Characterization of YopM^ΔNΔC^ and the YopM^ΔNΔC^-hPYD complex by SEC. YopM^ΔNΔC^ (**A,B**) or a 1:1 molar mixture of YopM ^ΔNΔC^ and hPYD (**C,D**) were analyzed by SEC on a Superdex 200 10/300 column at 0.75 mL/min. Samples were analyzed in a buffer containing 20 mM HEPES, 150 mM NaCl and 1 mM TCEP, pH 7.4. **A,C**) SEC chromatograms, with fraction numbers in blue above elution volume. **B,D**) Fractions of the peak elutions were analyzed by SDS-PAGE and blue protein staining.

**Fig. S8.**
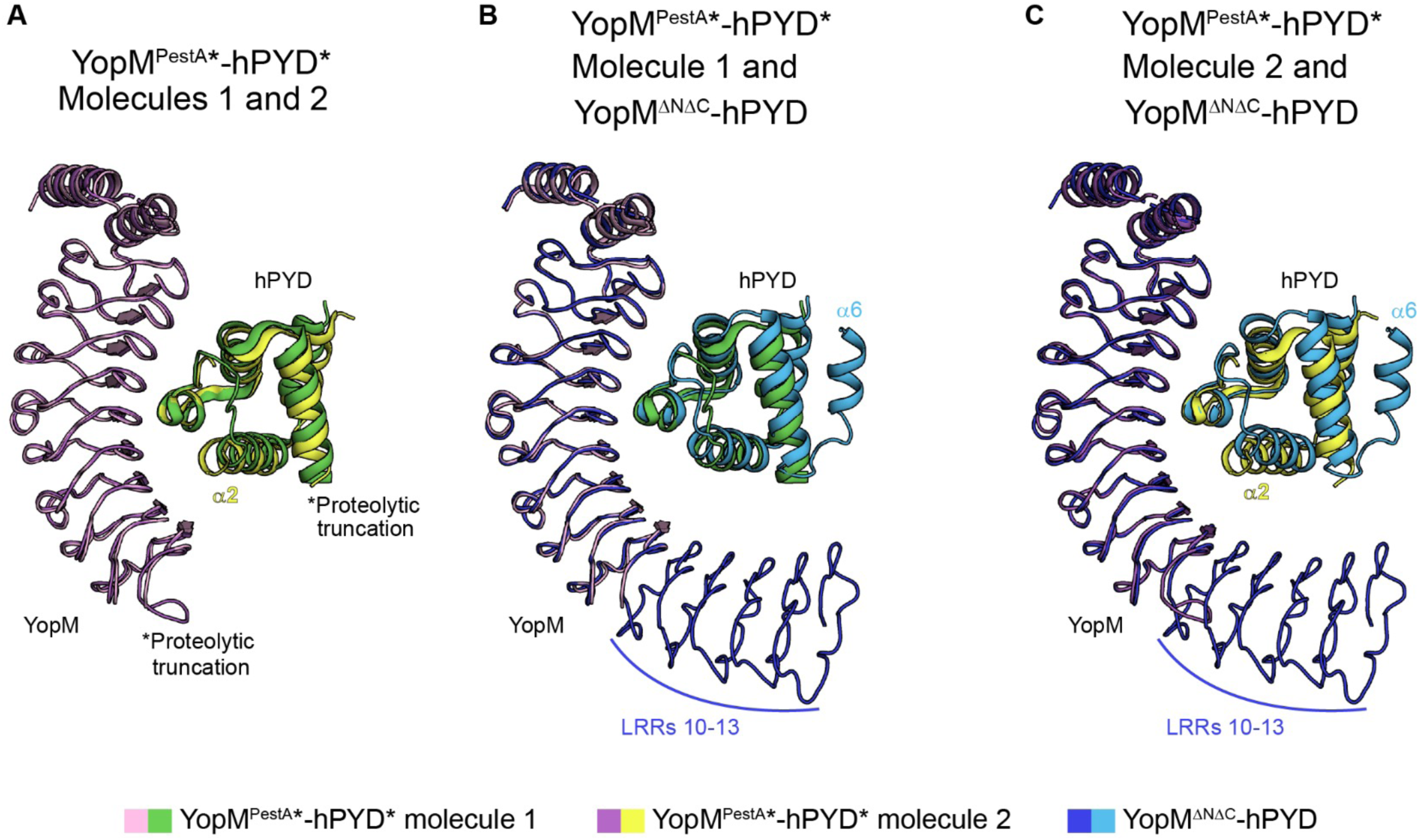
Heterogeneity in hPYD α2 is revealed by alignment of YopM^PestA^-hPYD complexes. The complexes of YopM^PestA^ (pink, deep purple, and dark blue) and hPYD (green, yellow, and light blue) were aligned by all C_a_ carbons in YopM^PestA^, and key structural differences are labeled. **A**) Alignment of both molecules in the ASU of the limited proteolysis structure (YopM^PestA^*-hPYD*) reveal differences in the position of hPYD* α2 where one molecule makes additional interactions with the concave surface of YopM^PestA^*. **B and C**) Additional alignments of each molecule in (A) to a second co-crystal structure of YopM^ΔNΔC^-hPYD where all YopM LRRs and hPYD α-helices were resolved. The positioning of hPYD α2 is consistent between YopM^PestA^*-hPYD* molecule 1 and YopM^ΔNΔC^-hPYD, as shown in (B).

**Fig. S9.**
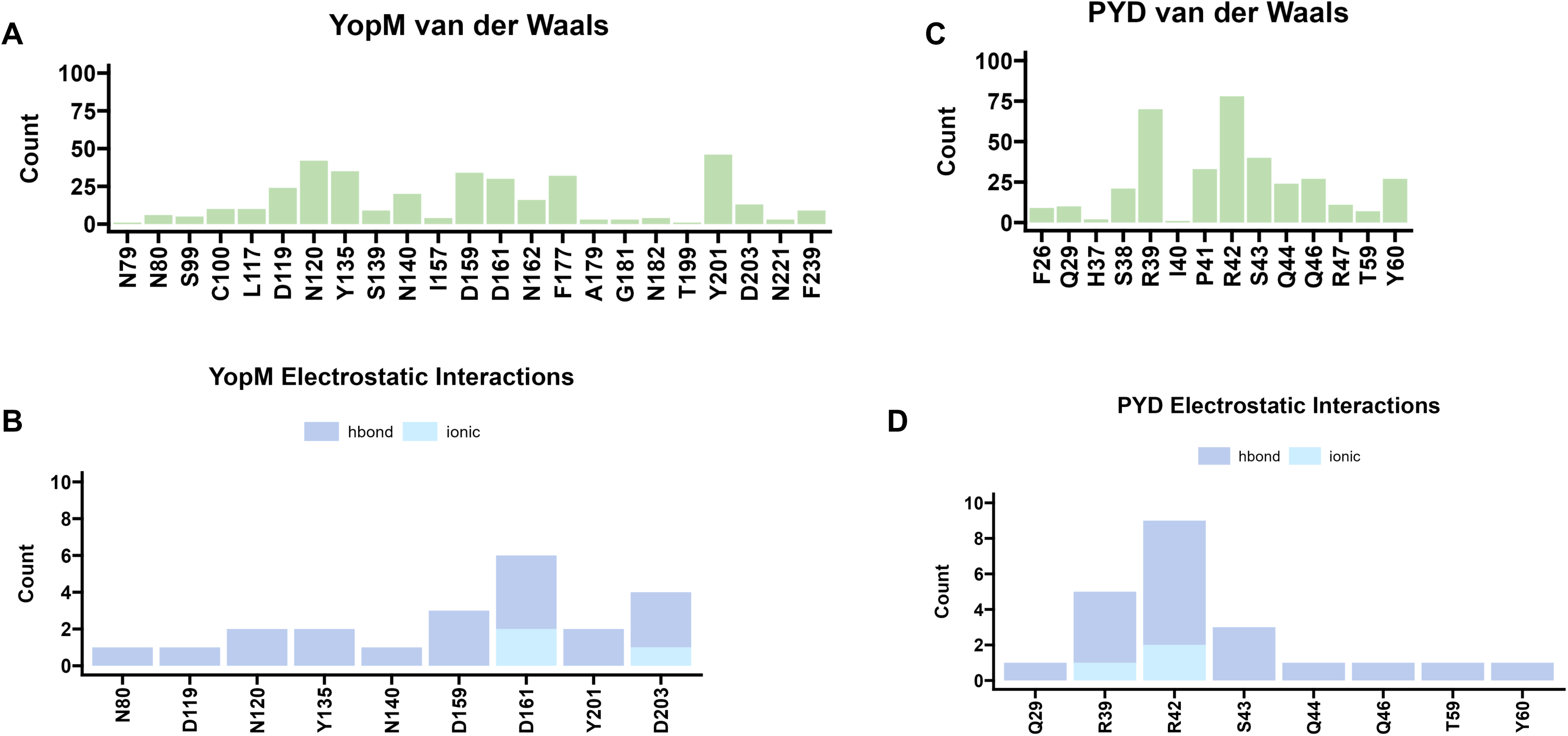
Residues contributing to the binding interface of YopM^PestA^-hPYD. The frequency with which each residue makes an interaction is plotted as counts on the y-axis. The y-axis denotes the residue position for the indicated protein**. A,C**) Plots of van der Waals interactions. **B,D**) Plots of electrostatic interactions showing hydrogen bonds in dark blue and ionic charges in light blue.

**Fig. S10.**
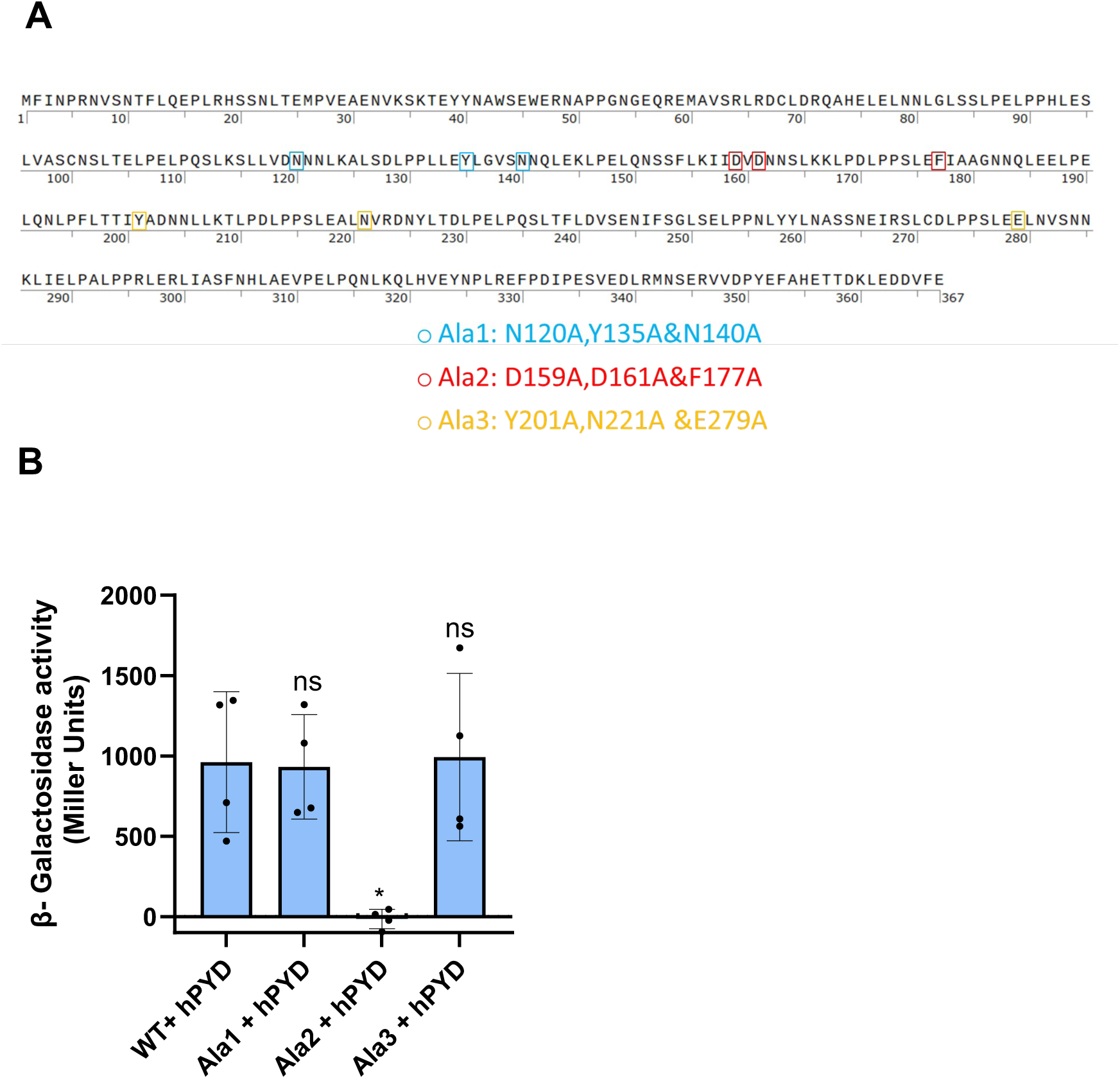
Identification by Ala substitution of amino acids in YopM^PestA^ that are important for hPYD binding. **A**) Amino acid sequence of YopM^PestA^ highlighting residues in the three-series alanine substitution variants by color: Ala1 (blue), Ala2 (red) and Ala3 (orange). **B**) Analysis of YopM^PestA^ Ala variants binding to hPYD using BACTH. Plasmids with the indicated fused partners were co-transformed into *E. coli* to test for protein-protein interactions as in Fig.1. Results are pooled from three independent experiments. Significance determined by one-way ANOVA and Tukey’s post hoc test comparing to PestA WT + hPYD; *p<0.05; ns, not significant.

**Fig. S11.**
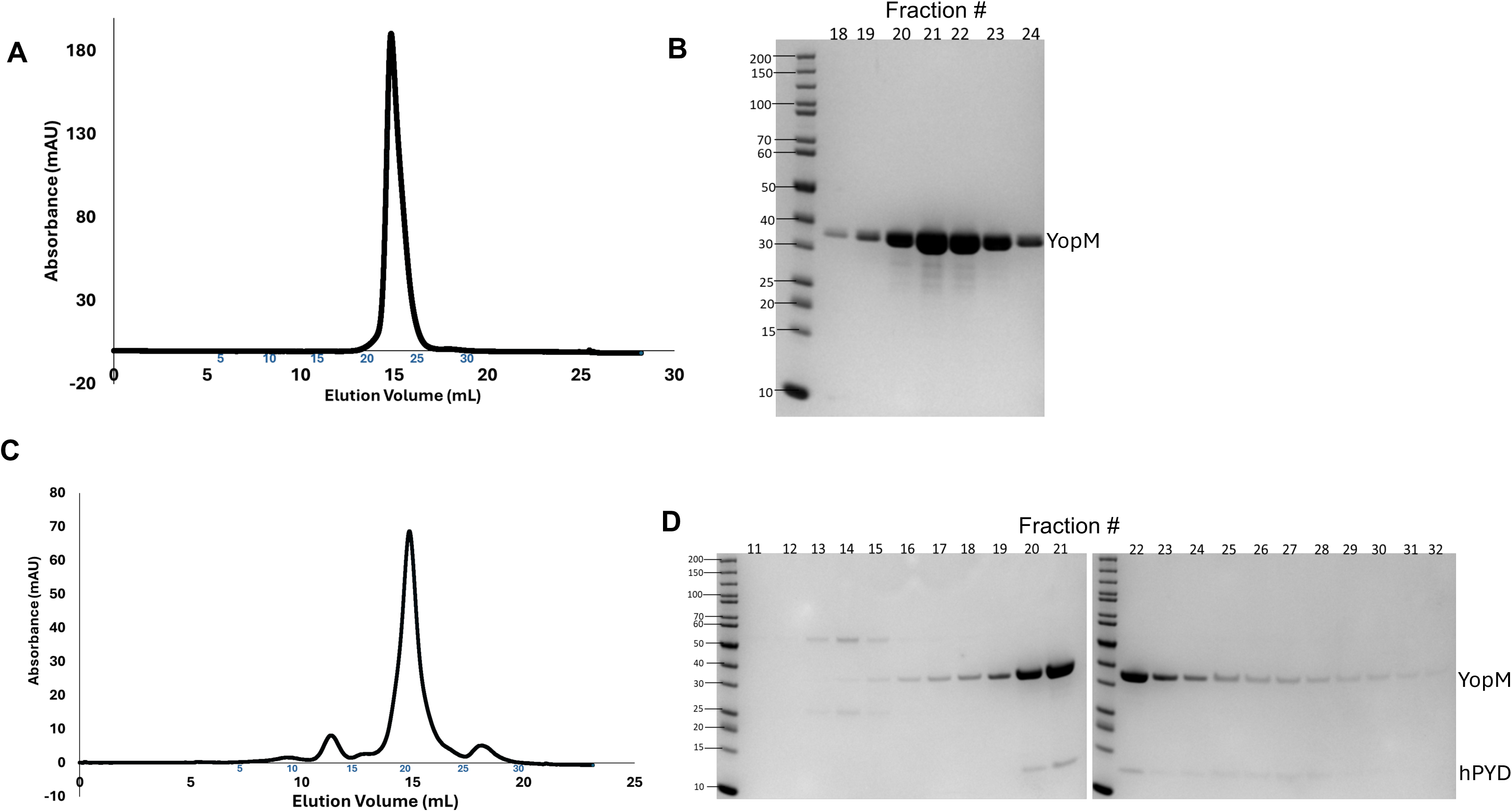
Characterization of YopM^ΔNΔC^ Ala2 and the YopM^ΔNΔC^ Ala2-hPYD interaction by SEC. YopM^ΔNΔC^ at 4mg/mL (**A,B**) or a 1:1 molar mixture of YopM ^ΔNΔC^ and hPYD at 2.3mg/mL (**C,D**) were analyzed by SEC on a Superdex 200 10/300 column at 0.75mL/min. Samples were analyzed in 20mM HEPES, 150mM NaCl and 1mM TCEP pH 7.4 buffer. **A,C**) SEC chromatograms. **B,D**) Fractions of the peak elutions were analyzed by SDS-PAGE and blue protein staining.

**Fig. S12.**
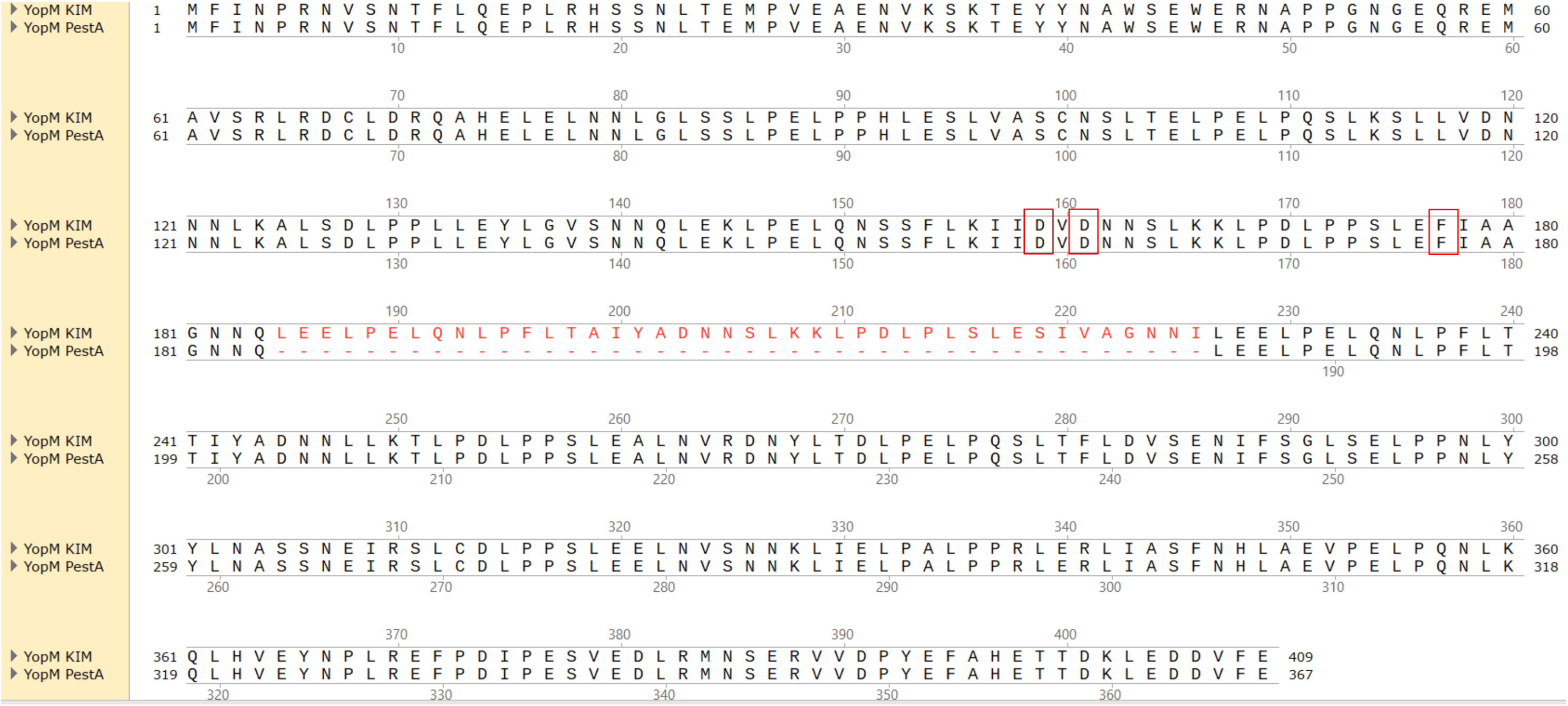
Sequence alignment of YopM^KIM^ and YopM^PestA^. Full-length sequences of the two proteins were aligned in Snapgene using ClustalW. Highlighted in red boxes are the residues changed in Ala2. In red text are the residues in YopM^KIM^ which are missing in YopM^PestA^ (red dashes).

**Fig. S13.**
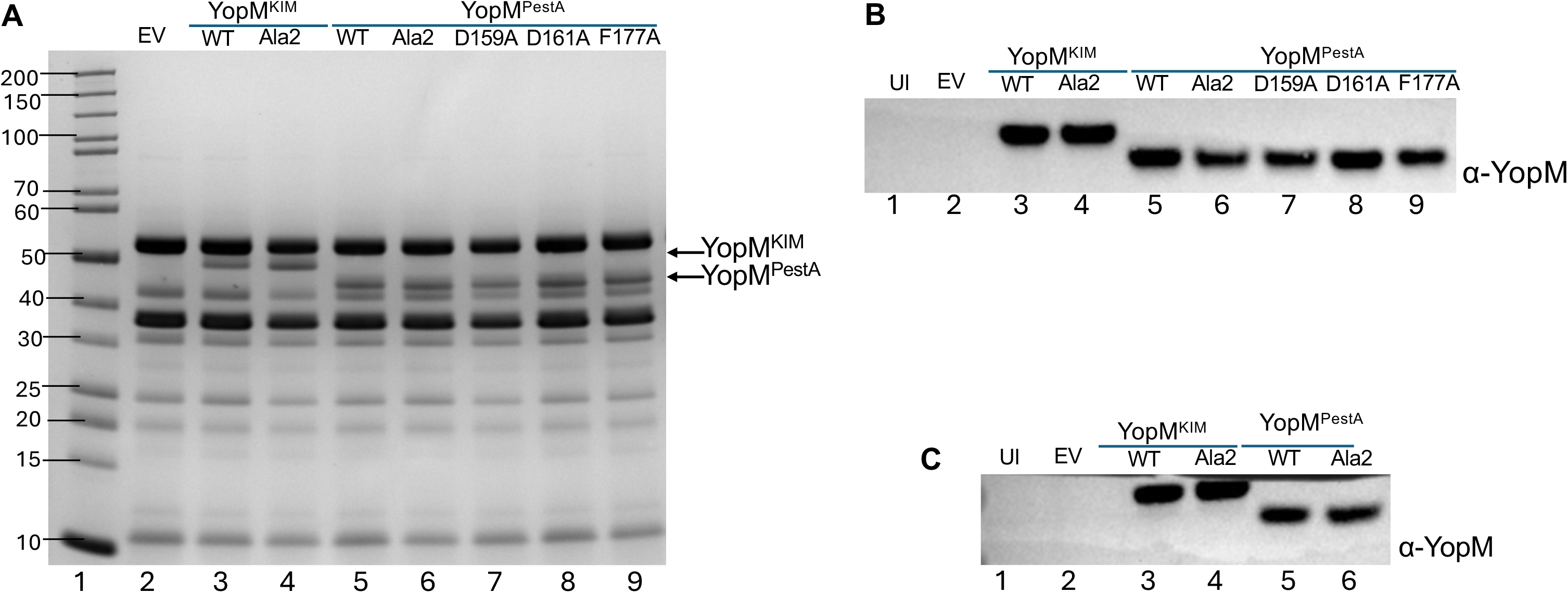
Analysis of YopM variant secretion and translocation. **A**) *Y. pseudotuberculosis yopJ^C172A^ΔyopM* strains complemented with empty vector (EV) or expressing the indicated YopM isoform or variant were grown under T3SS-inducing conditions and secreted Yops were collected and analyzed by SDS-PAGE and blue protein staining. THP-1 cells (**B**) or BMDMs (**C**) were left uninfected (UI) or infected with *Y. pseudotuberculosis yopJ^C172A^ΔyopM* strains complemented with empty vector (EV) or expressing the indicated YopM isoform or variant. THP-1 infections were as per Figs. 4C and 4D. BMDM infections were at MOI 30 for 90 min. Cell lysates were subjected to Immunoblotting with α-YopM.

**Fig. S14.**
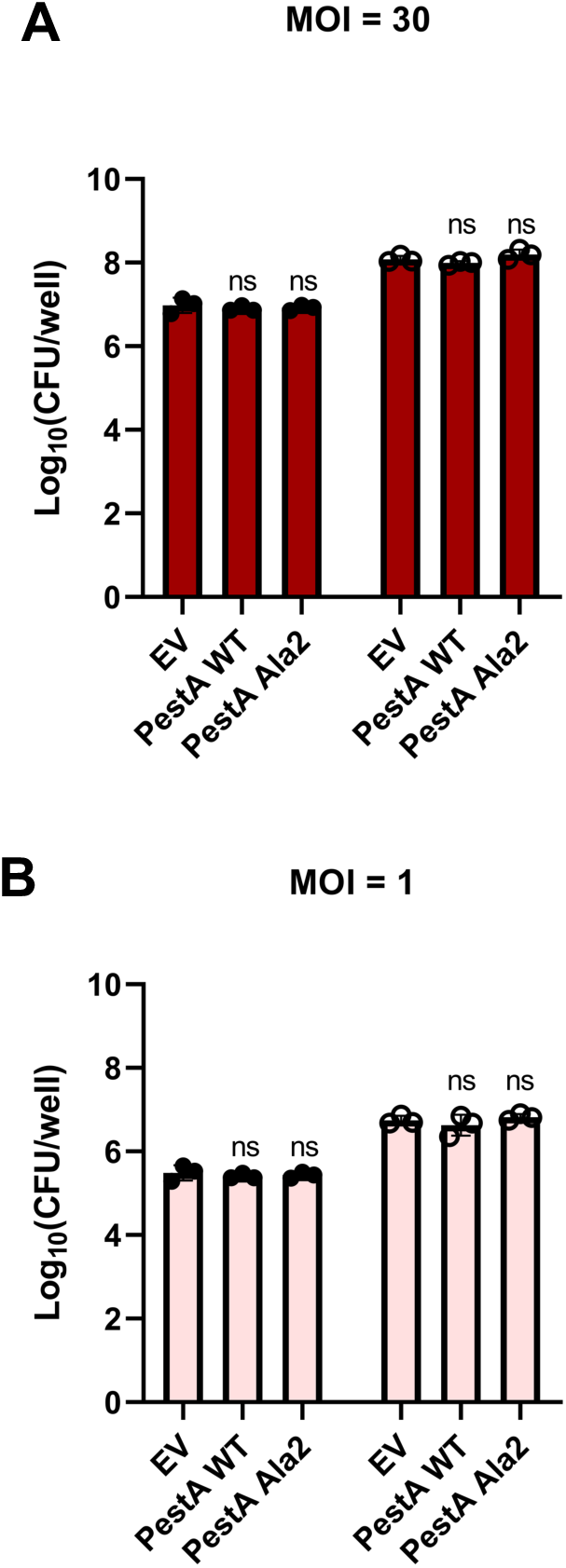
CFU recovery post-infection. THP-1 cells were infected with the indicated strains of *Y. pseudotuberculosis* at an MOI=30 (**A**) or MOI=1 (**B**) for 4 h. Samples of bacteria inoculated into each well at t=0 h (closed circles) were serially diluted and plated for CFU quantification. Total bacterial-monocyte co-cultures were solubilized in 0.1% NP-40 detergent (t=4 h) followed by serial dilutions and plating to determine CFU recovery post-infection (open circles). Bars represent averages of 3 biological replicates. Error bars represent standard deviation. Significance was determined by one-way ANOVA and Tukey’s *post hoc* test. Comparisons of each group to EV are shown; ns, not significant.

**Fig. S15.**
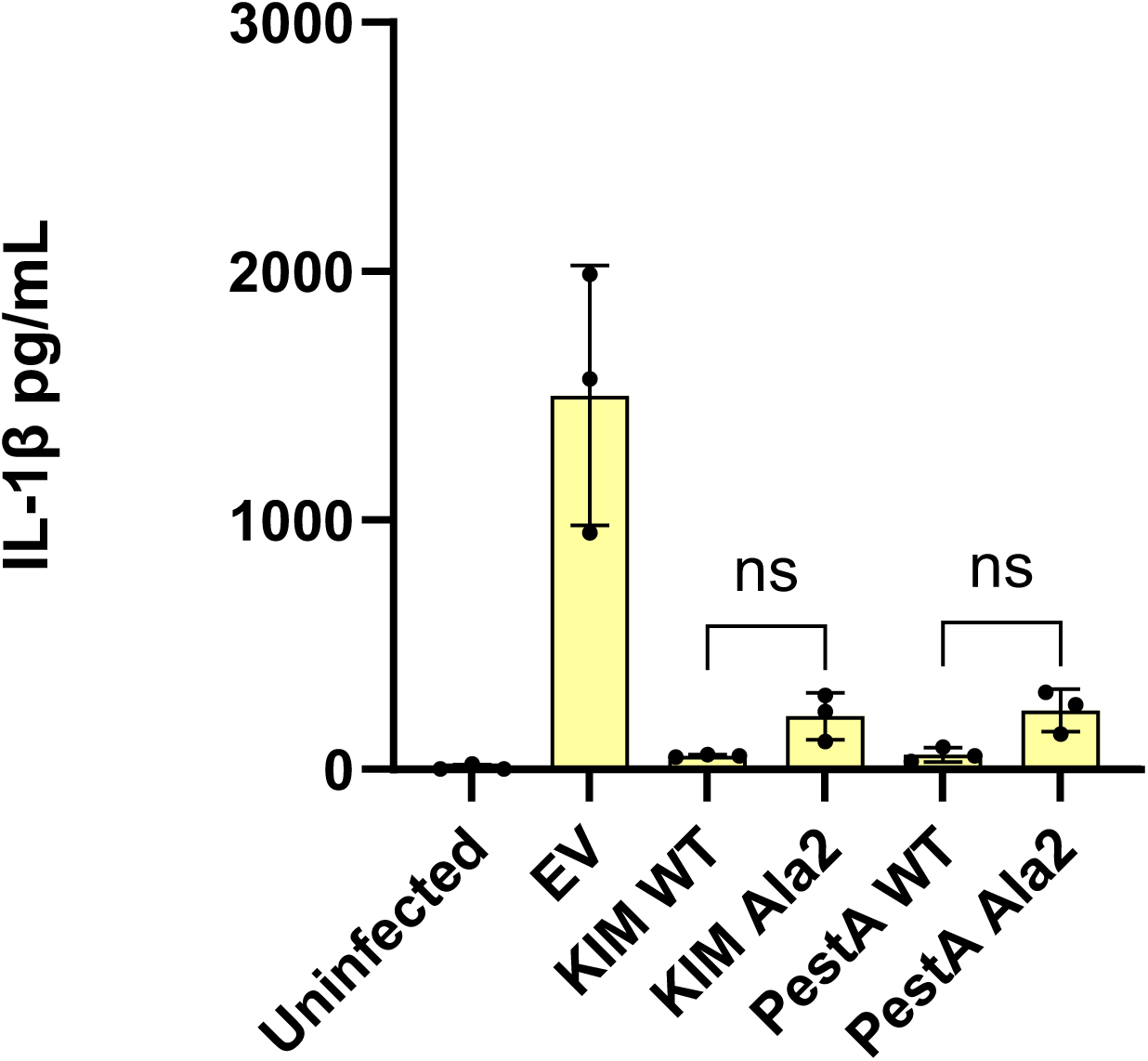
Analysis of YopM variants by BMDM infection assay. LPS-primed BMDMs were left uninfected (UI) or infected with *Y. pseudotuberculosis yopJ^C172A^λ1yopM* strains complemented with empty vector (EV) or expressing the indicated YopM isoform or variant. ELISA was used to measure released IL-1β. Results are averages and standard deviations with data pooled from at least 3 independent experiments. Significance determined by one-way ANOVA and Tukey’s post hoc test comparing WT YopM isoforms to their Ala2 variants as shown by brackets. ns, not significant.

**Fig. S16.**
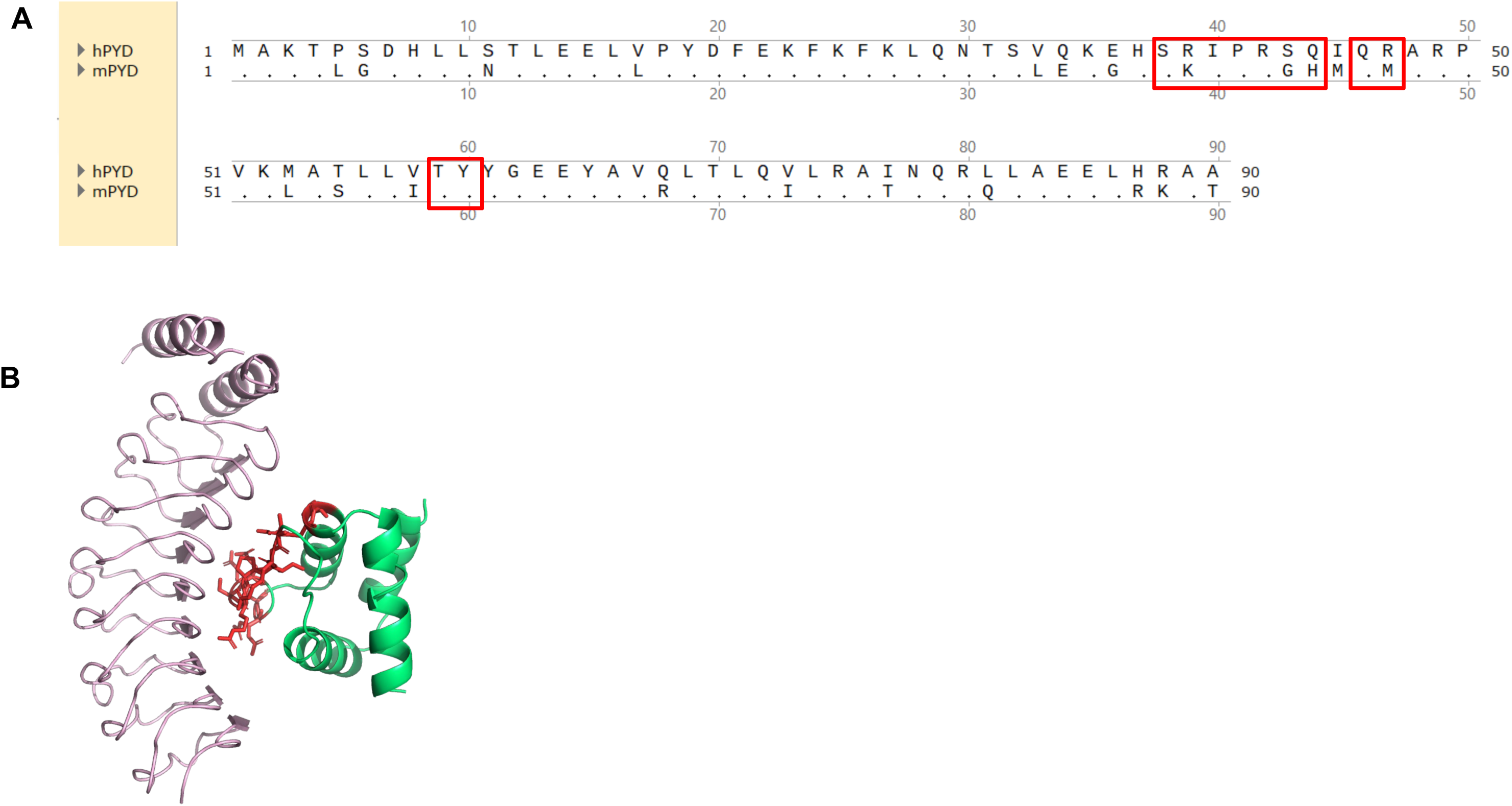
Sequence alignment of hPYD and mPYD and structure of the complex highlighting key contact residues in hPYD. **A**) Sequences of hPYD and mPYD were aligned in Snapgene using ClustalW. Dots indicate identical residues at each position. Highlighted in red boxes are residues on hPYD participating in key interactions with YopM. **B**) YopM*-hPYD* complex highlighting residues on hPYD* as sticks in red that participate in key interactions with YopM*.

